# Characterization of an early-diverging KCNE potassium-channel auxiliary subunit in the jawless vertebrate lamprey

**DOI:** 10.64898/2026.04.28.721385

**Authors:** Go Kasuya, Kaei Ryu, Buntaro Zempo, Emi Kawano-Yamashita, Koichi Nakajo

## Abstract

The KCNE (KCNE1–6) proteins are single-pass transmembrane auxiliary subunits of the voltage-gated K^+^ channel KCNQ1. KCNQ1–KCNE complexes have been well studied in jawed vertebrates ranging from zebrafish to humans, but KCNE subunits from earlier-diverging vertebrates remain poorly characterized. Here, we functionally characterize a single KCNE-like gene in lamprey, a jawless vertebrate, and designate it *kcne0* as an early-diverging member of the KCNE family. KCNE0 shows moderate amino acid sequence similarity to KCNE1–6 but is not particularly similar to any single isoform. Both *kcnq1* and *kcne0* transcripts were detected in multiple lamprey organs. When co-expressed with lamprey KCNQ1, KCNE0 produced a constitutively active current, similar to KCNE3. By contrast, KCNE0 modulated KCNQ1 from other species less effectively, suggesting species-specific tuning of KCNQ1–KCNE compatibility. Introducing into KCNE0 an intracellular tetra-leucine motif analogous to that in KCNE4 markedly reduced KCNQ1 current amplitude, conferring a KCNE4-like inhibitory effect. Overall, this work provides a functional reference for comparing KCNE-dependent modulation of KCNQ1 across vertebrates and suggests an underlying compatibility mechanism.

## Introduction

Modulation of ion channels by auxiliary subunits is essential for generating diverse physiological functions^1,2^. KCNE proteins are single-pass transmembrane auxiliary subunits of the voltage-gated K⁺ channel KCNQ1 (Kv7.1). In jawed vertebrates, six members (KCNE1–6) have been identified^3–10^. Among them, KCNE1 and KCNE3 are the best studied because they produce opposite effects on KCNQ1 gating with clear physiological roles. KCNE1 shifts the voltage dependence of KCNQ1 to more positive potentials to generate the slow delayed-rectifier K⁺ current (I_Ks_), which is essential for ventricular repolarization and inner-ear K⁺ homeostasis^11–14^. By contrast, KCNE3 shifts the voltage dependence to more negative potentials to produce a constitutively active current that supports epithelial K⁺ recycling^1,2,7^. Other KCNE subunits also modulate KCNQ1 gating, but their physiological roles are not well understood. For example, KCNE2 produces a constitutively active current, whereas KCNE5 and KCNE6 shift it to more positive potentials. KCNE4 suppresses KCNQ1 current ^2,10,15,16^.

Since the first KCNE member was cloned from rat kidney (originally described as minK; rat *Kcne1*)^3^, KCNE subunits have been identified and characterized not only in humans but also in a variety of non-human jawed vertebrates, including mice^7,17^, horses^18^, frogs^19^, and zebrafish^20,21^. Comparative genomic analyses have not identified canonical KCNE genes in invertebrates^22^, although *Caenorhabditis elegans* encodes *mps1–4*, nematode-specific single-pass membrane proteins that are sometimes described as MiRP/KCNE-like^23,24^. Together, these observations suggest that canonical KCNE genes expanded and became widespread after the rise of jawed vertebrates^22^. However, KCNE subunits from early-diverging vertebrates remain functionally uncharacterized, limiting our understanding of how KCNE-dependent KCNQ1 modulation originated and diversified.

To address this gap, we searched for KCNE-like sequences outside jawed vertebrates and found a single *KCNE*-like gene in lamprey, a cyclostome (jawless vertebrate) that diverged early in the vertebrate lineage^25,26^. RT-PCR and re-analysis of deposited RNA-seq data^27^ detected transcripts of *kcnq1* and the KCNE-like gene in multiple lamprey organs. When co-expressed with the lamprey KCNQ1, this lamprey KCNE-like subunit produced constitutive activity, similar to KCNE3, and modestly reduced current amplitude. By contrast, its effect on KCNQ1 from other species was weaker, suggesting species-specific compatibility between KCNQ1 and KCNE. Introducing into the lamprey subunit a short intracellular tetra-leucine motif homologous to that in human KCNE4 markedly reduced KCNQ1 current, conferring a KCNE4-like inhibitory effect. Based on these sequences and functional data, we designate this subunit KCNE0 and propose that it represents an early-diverging member of the KCNE family.

## Results

### Characterization of lamprey KCNE0

Guided by previous comparative analyses that have not identified canonical KCNE family genes in invertebrates^22^, we surveyed cyclostomes, the extant jawless vertebrates including lampreys and hagfishes, as an early-diverging vertebrate group^25,26^. Public genome databases at NCBI and Ensembl revealed no annotated *kcne*-like sequences in hagfish assemblies, whereas a single *kcne*-like gene was present in two lamprey species, sea lamprey (*Petromyzon marinus*, Pm) and Far Eastern brook lamprey (*Lethenteron reissneri*, Lr), together with *kcnq1*. We additionally obtained Arctic lamprey (*Lethenteron camtschaticum*, Lc) and cloned its *kcne* and *kcnq1* cDNAs using the sea lamprey and Far Eastern brook lamprey sequences as templates (Supplementary Figs. 1 and 2). The three lamprey KCNE proteins share ∼95% amino-acid identity (Fig. 1A–C). In the Ensembl genome browser, the sea lamprey *kcne* locus lies near the *rab20* locus (Fig. 1D). Pairwise alignments against human KCNE1–5 and zebrafish KCNE6 showed moderate identity but did not unambiguously match a single isoform (Fig. 1A–C). Therefore, we refer to the lamprey subunit as KCNE0 throughout for clarity.

**Figure 1.**
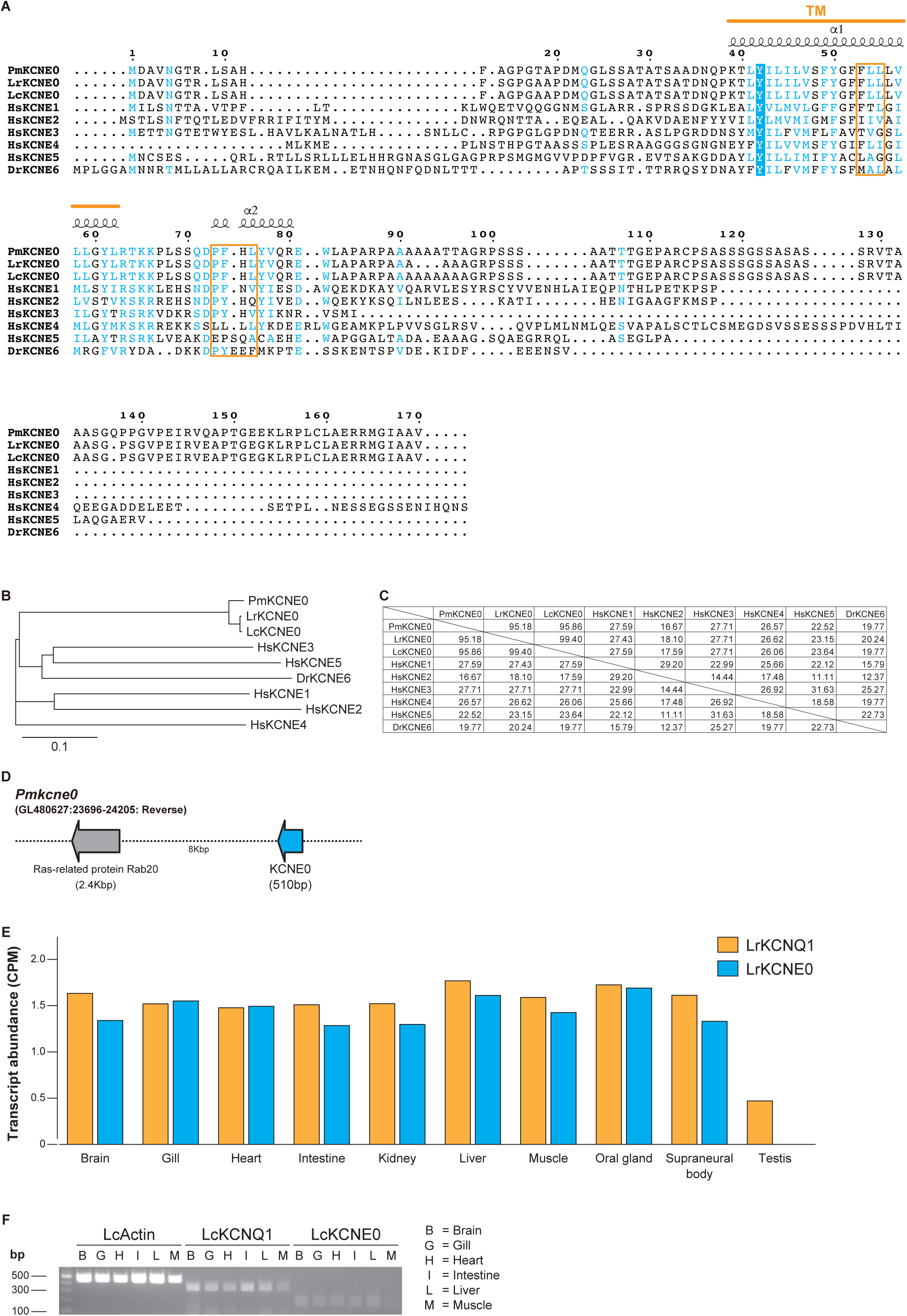
Sequence, genomic context, and transcript distribution of lamprey KCNE0. (**A–C**) Amino-acid sequence alignment (A), phylogenetic tree (B), and percent identity (C) of lamprey and human KCNE subunits. Alignments were generated with Clustal Omega^61^ and displayed using ESPript3^62^. Residues corresponding to the “triplet”^42,43^ motif are highlighted with an orange square. (**D**) Genomic region containing the *kcne* locus in sea lamprey, shown from the Ensembl genome browser^63^. For sequence comparison, KCNE0 from lamprey species (sea lamprey, PmKCNE0; NCBI Accession Number XP_032831116.1; Far Eastern brook lamprey, LrKCNE0; XP_061431545.1 and Arctic lamprey, LcKCNE0; see Supplementary Fig. 2), the five human KCNE subunits (HsKCNE1, NP_000210.2; HsKCNE2, NP_751951.1; HsKCNE3, NP_005463.1; HsKCNE4, NP_542402.4; and HsKCNE5, NP_036414.1), and zebrafish KCNE6 (DrKCNE6)^10^ were used. (**E**) RNA-seq-based transcript levels of KCNQ1 (XM_061564454.1) and KCNE0 (XM_061575561.1) from 10 organs of Far Eastern brook lamprey quantified using a decoy-aware Salmon index. Values are shown as counts per million of the library (CPM). (F) RT-PCR detection of KCNQ1 and KCNE0 transcripts using cDNA synthesized from total RNA isolated from six organs of Arctic lamprey.

To explore whether *kcne0* is co-expressed with *kcnq1* at the transcript level across lamprey tissues, we next examined the organ distribution of *kcnq1* and *kcne0* transcripts in lamprey. We analyzed deposited RNA-seq data from Far Eastern brook lamprey (*Lethenteron reissneri)* (BioProject PRJNA558325; SRX6711574–SRX6711583) covering 10 organs^27^ and quantified transcript levels. Transcripts of *kcnq1* and *kcne0* were detected in most tissues at low levels (approximately 1–1.5 counts per million, CPM), whereas testis showed lower *kcnq1* levels and *kcne0* was not detected under our analysis threshold (Fig. 1E). We next performed RT-PCR on cDNA synthesized from total RNA isolated from six organs of Arctic lamprey (*Lethenteron camtschaticum*) and detected amplicons for both *kcnq1* and *kcne0* (Fig. 1F and Supplementary Fig. 3). Although these data do not demonstrate protein expression or native complex formation, they indicate that *kcnq1* and *kcne0* transcripts show a broadly overlapping organ distribution in at least two lamprey species.

To investigate how KCNE0 modulates KCNQ1 gating in lamprey, we used two-electrode voltage clamp (TEVC) in *Xenopus laevis* oocytes. When expressed alone, all three lamprey KCNQ1 channels (PmKCNQ1 WT, LrKCNQ1 WT, and LcKCNQ1 WT) showed voltage-dependent activation with a sigmoidal conductance–voltage (G–V) relationship, comparable to KCNQ1 from other species. Their *V_1/2_* values differed modestly but significantly, with PmKCNQ1 activating at slightly more negative voltages than LrKCNQ1 and LcKCNQ1 (PmKCNQ1 WT, −35.8 ± 0.9 mV; LrKCNQ1 WT, −32.9 ± 0.6 mV; LcKCNQ1 WT, −32.1 ± 0.6 mV; n = 5 each; Supplementary Table 1). By contrast, co-expression with their corresponding KCNE0 WT subunits produced a constitutively active current, resembling the mammalian KCNQ1–KCNE3 complex (Fig. 2A–G; Supplementary Table 1).

**Figure 2.**
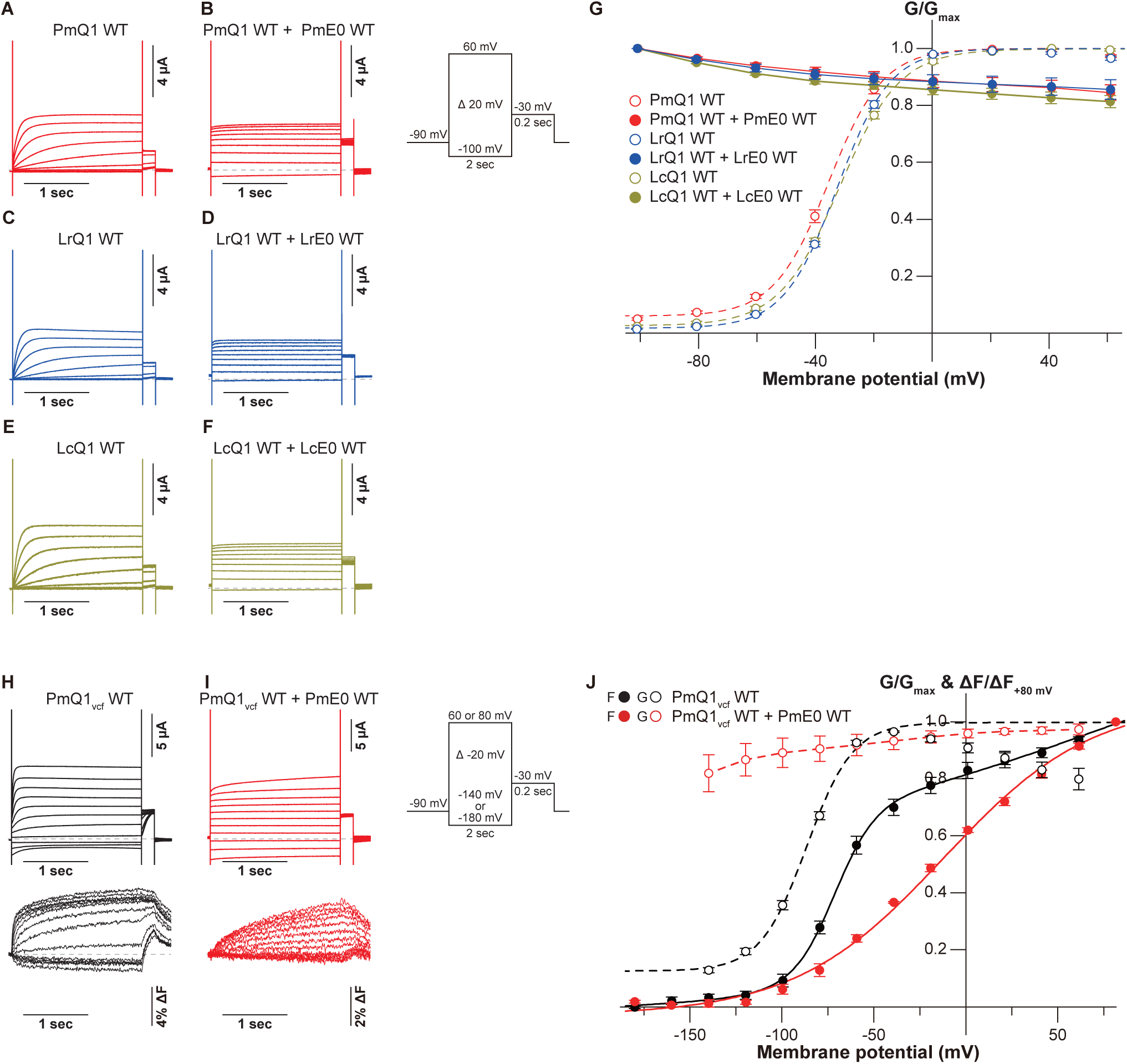
Biophysical properties of lamprey KCNQ1 and modulation by KCNE0. (**A–G**) Representative current traces (A–F) and G–V relationships of lamprey KCNQ1 WT expressed alone or co-expressed with the corresponding KCNE0 WT. (**H–J**) Representative ionic-current and fluorescence traces (H,I) and F–V relationship (J) of PmKCNQ1_vcf_ WT expressed alone or co-expressed with PmKCNE0 WT. Error bars denote mean ± s.e.m. for n = 5 in (G,J). Statistical comparisons of G–V *V_1/2_* values for lamprey KCNQ1 WT expressed alone are provided in Supplementary Table 1. *V_1/2_* values were not determined for KCNQ1–KCNE0 complexes because the G–V relationships were nearly saturated over the tested voltage range.

To examine whether the KCNE0-dependent G–V changes arise from altered voltage-sensor movements, we performed voltage-clamp fluorometry (VCF). VCF monitors fluorescence from a dye attached near the fourth transmembrane segment (S4), the principal voltage sensor of KCNQ1, so changes in the fluorescence–voltage (F–V) relationship report conformational movements of the voltage-sensor domain (VSD) and allow direct comparison of F–V and G–V relationships in the same construct^28^. Because the PmKCNQ1/PmKCNE0 pair produced the most consistent recordings, we focused on this pair for VCF and subsequent biophysical analyses. We generated a VCF construct (PmKCNQ1 C205A/G210C; PmKCNQ1_vcf_ WT) by introducing a cysteine mutation at position G210 in the extracellular S3–S4 loop, aligned with the site commonly used for VCF in human KCNQ1 (G219)^29,30^ (Supplementary Fig. 4). Previous VCF studies of human KCNQ1 have shown that KCNQ1 expressed alone often exhibits an F–V relationship that is well described by a double-Boltzmann function, consistent with stepwise VSD activation. Within the established six-state gating framework of KCNQ1, channel gating is governed by three VSD positions (down, intermediate, and up/activated) and two pore domain (PD) conformations (closed and open). Accordingly, KCNQ1 can open in an intermediate-open (IO) and an activated-open (AO) state during VSD activation^31–33^. In such recordings, the dominant fluorescence component (F_1_), corresponding to the transition from down to intermediate VSD positions, typically overlaps with the G–V relationship.

In our analysis, PmKCNQ1_vcf_ WT expressed alone showed overall features similar to human KCNQ1. Its G–V relationship was fit with a single-Boltzmann function, whereas the F–V relationship was fit with a double-Boltzmann function. The dominant F_1_ overlapped with the G–V relationship, consistent with channel opening within the established IO/AO framework^31–33^ (Fig. 2H–J; Supplementary Table 1). By contrast, co-expression of PmKCNE0 altered the fluorescence behavior of PmKCNQ1_vcf_ WT. The G–V relationship remained close to 1 over the tested voltage range, whereas the F–V relationship was best described by an apparent single-Boltzmann function. This suggests that the down-to-intermediate VSD transition occurs at voltages more negative than those accessible in our recordings, such that the corresponding fluorescence component was not resolved. Accordingly, the observed F–V relationship indicates stabilization of an intermediate-like VSD conformation (Fig. 2H–J; Supplementary Table 1). Consistent with this interpretation, similar KCNE-dependent changes in KCNQ1 VSD motion have been widely reported, in which KCNE subunits alter VSD-PD coupling and stabilize distinct VSD positions. For example, KCNE1 stabilizes an intermediate closed (IC) state, whereas KCNE3 stabilizes an intermediate open (IO) state at most physiological voltage ranges^31,33–39^. Taken together, these results indicate that lamprey KCNE0 modulates KCNQ1 gating, at least in part, by altering VSD movement, consistent with mechanisms described for the mammalian KCNQ1–KCNE3 complex.

Next, to identify which regions of KCNE0 are required for modulating KCNQ1 gating, we generated a series of PmKCNE0 truncations at the N and C termini and co-expressed them with PmKCNQ1 WT (Fig. 3A). For N-terminal truncations, removing as few as six residues (PmKCNE0ΔN6) largely abolished modulation of PmKCNQ1 (Fig. 3B–F; Supplementary Table 1). For C-terminal truncations, the modulatory effect decreased progressively as the C terminus was shortened and was nearly lost in the construct lacking 105 residues (PmKCNE0ΔC105) (Fig. 3G–N; Supplementary Table 1). Confocal imaging indicated that neither N- nor C-terminal truncations detectably altered membrane localization under our conditions (Fig. 3O–T). Overall, the KCNE0 N-terminus is highly sensitive to truncation for proper modulation of KCNQ1, while the C-terminus is more tolerant and loses most of its effect only after deleting 105 residues (ΔC105).

**Figure 3.**
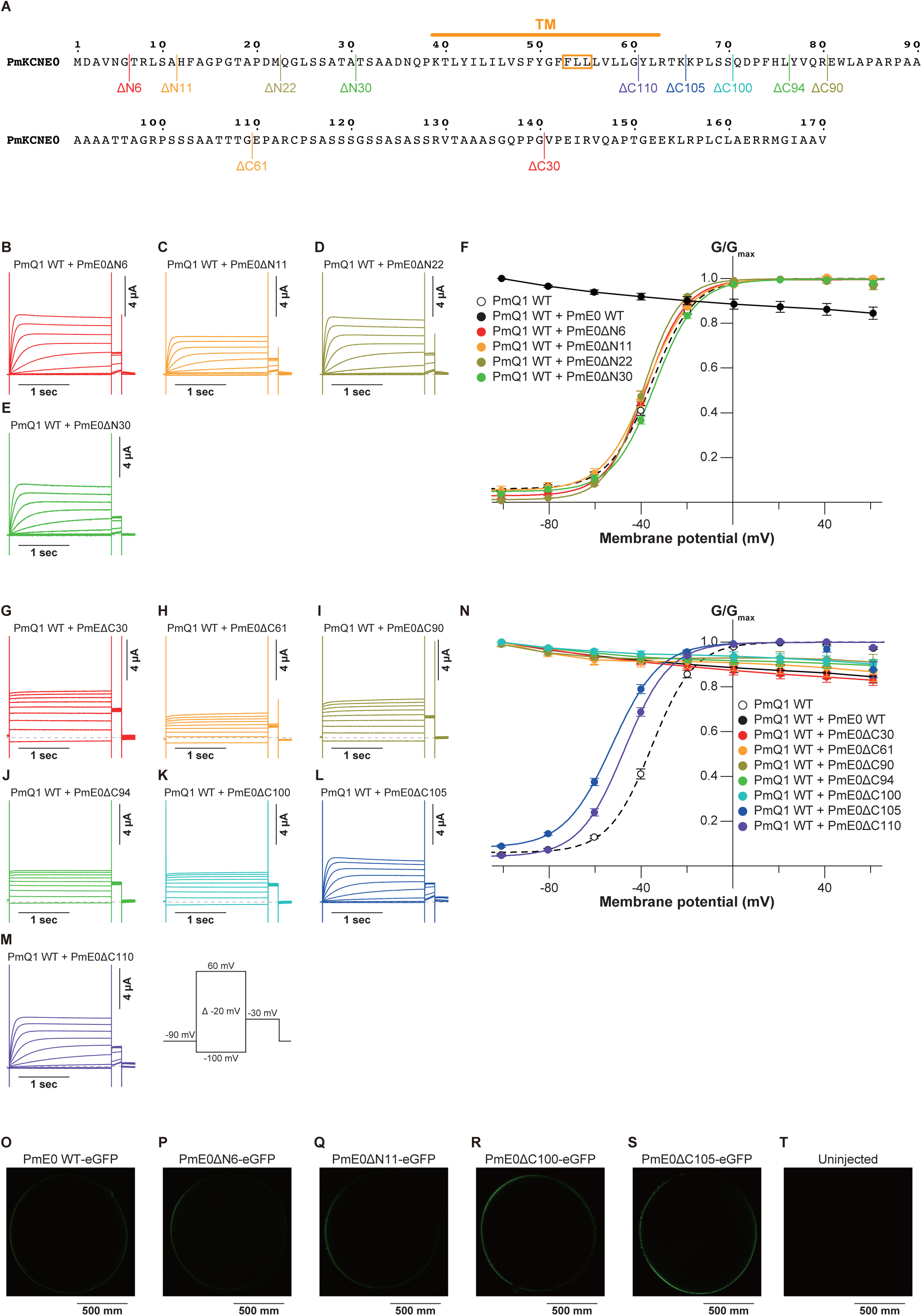
Regions of KCNE0 important for KCNQ1 modulation. (**A**) Schematic diagram of N- and C-terminal truncation constructs of PmKCNE0. (**B–F**) Representative current traces (B–E) and G–V relationship (F) of PmKCNQ1 WT expressed alone or co-expressed with each of the N-terminally truncated PmKCNE0 constructs. (**G–N**) Representative current traces (G–M) and G–V relationship (N) of PmKCNQ1 WT expressed alone or co-expressed with each of the C-terminally truncated PmKCNE0 constructs. Error bars denote mean ± s.e.m. for n = 5 in (F,N). (**O–T**) Confocal images of oocytes expressing C-terminally eGFP-tagged PmKCNE0 WT or truncation constructs, and an uninjected oocyte (control).

### Species-specific tuning of KCNQ1–KCNE compatibility

Species-specific tuning of KCNQ1–KCNE interactions has been observed across chordates. For example, we previously reported that KCNQ1 from the vase tunicate *Ciona intestinalis* (CiKCNQ1) is not effectively modulated by mammalian KCNE subunits, including KCNE1 and KCNE3^40^. These findings suggest that functional compatibility between KCNQ1 and KCNE subunits is tuned in a species-specific manner. To examine whether KCNE0 modulation also exhibits such species-specific tuning, we performed cross-pairing experiments. We co-expressed PmKCNE0 WT with KCNQ1 WTs from human (*Homo sapiens*, Hs), zebrafish (*Danio rerio*, Dr), and vase tunicate (*Ciona intestinalis*, Ci). PmKCNE0 modulated HsKCNQ1 and DrKCNQ1, but the effects were weaker and less consistent than those observed with PmKCNQ1. In HsKCNQ1, PmKCNE0 produced a mixed effect, with partial constitutive activity at negative voltages and a positive shift of the half-activation voltage (*V_1/2_*; HsKCNQ1 WT, −26.5 ± 0.8 mV; HsKCNQ1 WT–PmKCNE0 WT, 14.6 ± 1.5 mV; n = 5 each) (Fig. 4A–C; Supplementary Table 1). In DrKCNQ1, PmKCNE0 produced partial constitutive activity at negative voltages with only a mild change in *V_1/2_* (DrKCNQ1 WT, −44.2 ± 1.1 mV; DrKCNQ1 WT–PmKCNE0 WT, −40.4 ± 1.3 mV; n = 5 each) (Fig. 4D–F; Supplementary Table 1). By contrast, CiKCNQ1 was not detectably modulated by PmKCNE0 under our conditions (Fig. 4G–I; Supplementary Table 1).

**Figure 4.**
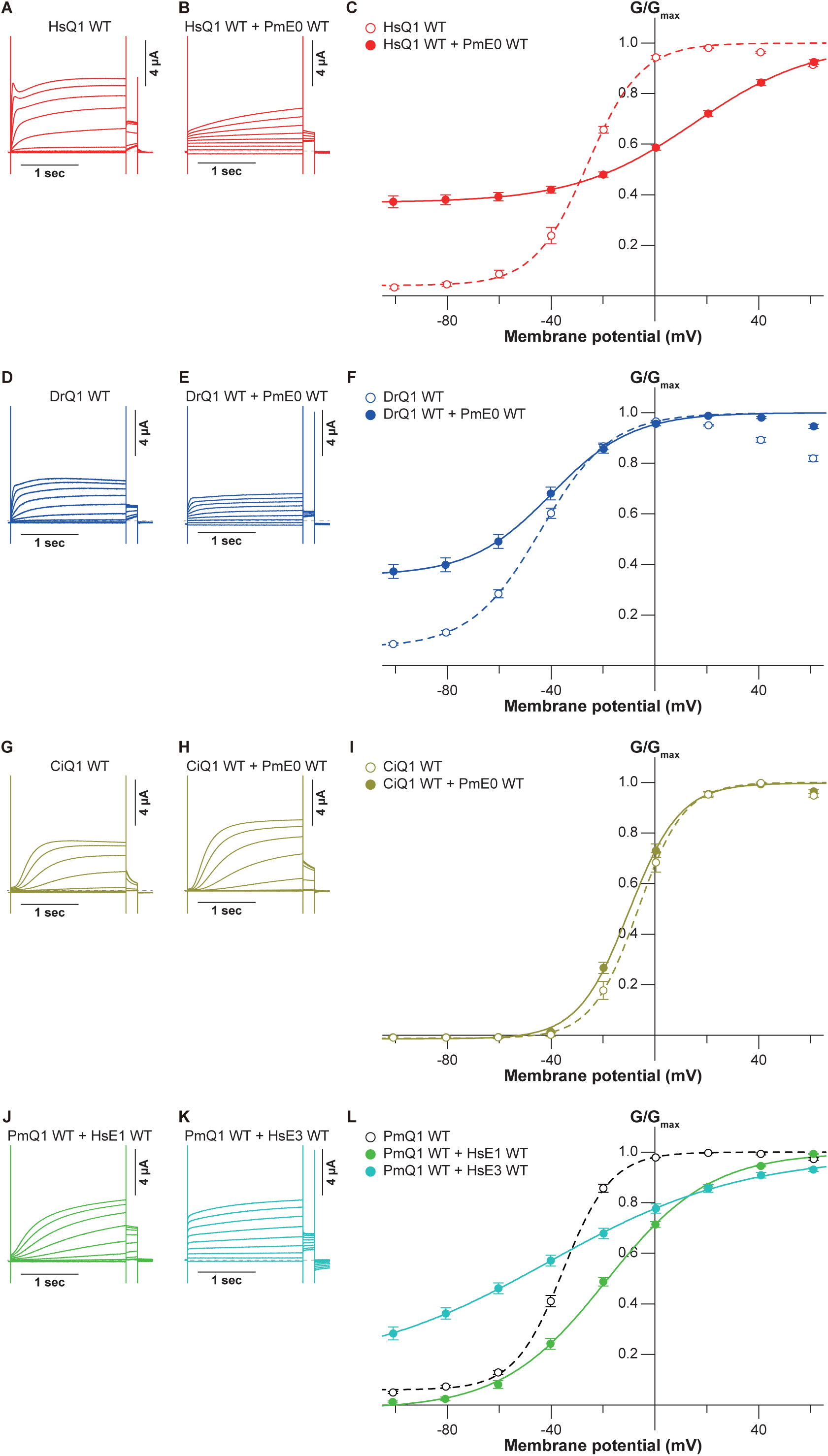
Species dependence of KCNQ1–KCNE compatibility. (**A–C**) Representative current traces (A,B) and G–V relationship (C) of HsKCNQ1 WT expressed alone or co-expressed with PmKCNE0 WT. (**D–F**) Representative current traces (D,E) and G–V relationship (F) of DrKCNQ1 WT expressed alone or co-expressed with PmKCNE0 WT. (**G–I**) Representative current traces (G,H) and G–V relationship (I) of CiKCNQ1 WT expressed alone or co-expressed with PmKCNE0 WT. (**J–L**) Representative current traces (J,K) and G–V relationship (L) of PmKCNQ1 WT co-expressed with HsKCNE1 WT or HsKCNE3 WT. Error bars denote mean ± s.e.m. for n = 5 in (C,F,I,L).

Reciprocally, we tested whether lamprey KCNQ1 is efficiently modulated by human KCNE subunits. Co-expression of PmKCNQ1 with HsKCNE1 produced only a small positive shift of *V_1/2_* (PmKCNQ1 WT, −35.8 ± 0.9 mV; PmKCNQ1 WT–HsKCNE1 WT, −19.3 ± 1.5 mV; n = 5 each) (Fig. 4J, L; Supplementary Table 1), which is markedly smaller than the strong positive shift typically observed for HsKCNQ1 with HsKCNE1 (about +50 mV positive shift)^1,3^. Likewise, pairing PmKCNQ1 with HsKCNE3 produced only a small negative shift of *V_1/2_* (PmKCNQ1 WT, same as above; PmKCNQ1 WT–HsKCNE3 WT, −41.3 ± 1.2 mV; n = 5) and partial constitutive activity at negative voltages (Fig. 4K, L; Supplementary Table 1), weaker than the robust constitutive effect typically observed for HsKCNQ1 with HsKCNE3^1,7^. Together, these cross-species results support the idea that KCNQ1–KCNE functional compatibility is tuned in a species-specific manner.

### Intracellular leucine substitutions shift KCNE0 toward a KCNE4-like inhibitory effect

Because KCNE subunits can show diverse effects on KCNQ1 gating^3–10^, we asked whether the early-diverging KCNE0 phenotype can be shifted by a small number of mutations. We first tested a KCNE4-related intracellular leucine motif that has been linked to KCNQ1 inhibition. KCNE4 contains a tetra-leucine sequence in a juxtamembrane intracellular region that is required for interaction with calmodulin and functional suppression of KCNQ1^41^ (Fig. 5A, B). Because PmKCNE0 already contains one leucine in the corresponding region (L76), we first added one leucine just before L76 by introducing H75L. This single substitution strongly reduced PmKCNQ1 current amplitude compared with PmKCNE0 WT, while the remaining current still showed constitutive activity across the tested voltages (Fig. 5C–F). We next introduced P73L, F74L, and H75L to create a tetra-leucine stretch from L73 to L76 together with the native L76. This change further attenuated KCNE0-dependent activation and shifted the G–V relationship toward that of PmKCNQ1 expressed alone (Fig. 5C–F; Supplementary Table 1). These results show that KCNE0 can be shifted toward a KCNE4-like inhibitory effect by introducing leucine substitutions in a small intracellular region. By contrast, we tested whether KCNE0 could be shifted toward a KCNE1-like effect. A previous study^37^ showed that the KCNE3 effect can be partially shifted toward a KCNE1-like effect by mutations in the middle of the KCNE transmembrane segment (the “triplet” motif)^42,43^ (Supplementary Fig. 5A, B; Supplementary Table 1). We therefore introduced an HsKCNE1-like change into the corresponding region of PmKCNE0 (PmKCNE0 L54T). However, this mutant produced a constitutively active current when co-expressed with PmKCNQ1, similar to PmKCNE0 WT (Supplementary Fig. 5C, D), indicating that a KCNE1-like modulatory effect cannot be readily recapitulated by introducing a single motif analogous to that of human KCNE1.

**Figure 5.**
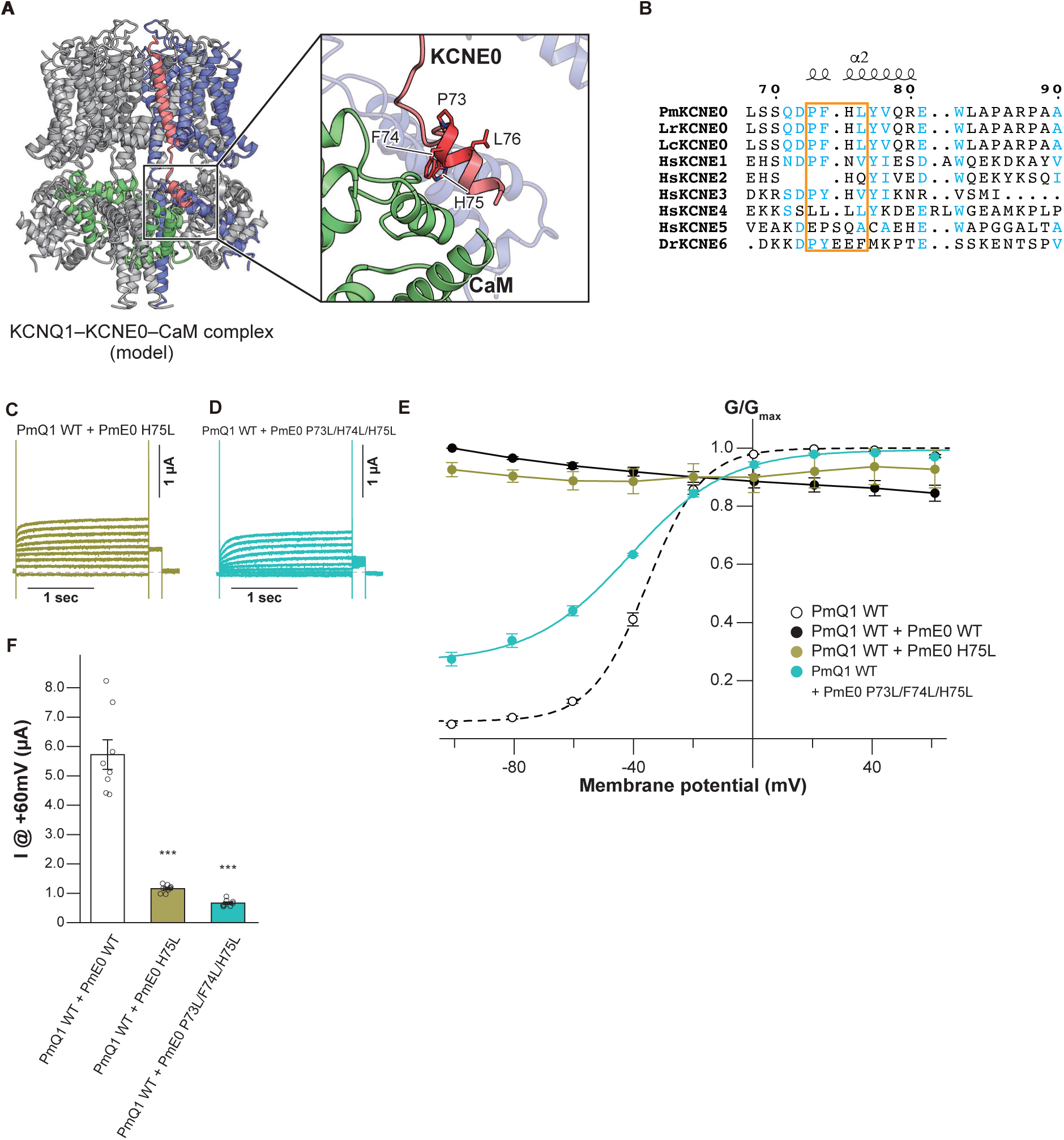
Intracellular leucine substitutions shift KCNE0 modulation toward a KCNE4-like inhibitory effect. (**A**) Close-up view of the interface between KCNQ1 and KCNE0 within the PmKCNQ1–PmKCNE0–PmCaM complex structure generated by SwissModel server^64^. The model was built with amino acid sequences of PmKCNQ1 (XP_075921450.1), PmKCNE0 (XP_032831116.1), and PmCaM (XP_032811771.1), using the human KCNQ1-KCNE3-CaM structure (PDB: 6V00) as a template. One KCNQ1, KCNE0 and CaM subunit are shown in blue, red, and green. The other regions are shown in gray. Residues used for motif-based mutagenesis are shown as sticks. Molecular graphics were prepared with CueMol (http://www.cuemol.org/). (**B**) Sequence alignment around the intracellular region corresponding to the KCNE4-related juxtamembrane tetra-leucine motif^41^. (**C–F**) Representative current traces (C,D), G–V relationship (E), and current amplitude at +60 mV (F) of PmKCNQ1 WT co-expressed with PmKCNE0 WT, PmKCNE0 H75L, or PmKCNE0 P73L/F74L/H75L. In (E), error bars denote mean ± s.e.m. for n = 5. In (F), error bars denote mean ± s.e.m. for n = 8. Statistical significance among the three constructs was assessed using one-way ANOVA followed by Tukey–Kramer multiple-comparison test. Significant differences are indicated by asterisks (***P < 0.001).

## Discussion

In this study, we functionally characterized a KCNE subunit from lamprey and designated it KCNE0. Because cyclostomes, the extant jawless vertebrates including lampreys and hagfishes, represent an early-diverging vertebrate lineage^25,26^, our results provide a useful reference point for understanding how KCNE-dependent modulation of KCNQ1 can operate outside jawed vertebrates. The lamprey KCNE subunit shows moderate amino acid sequence similarity to human KCNE1–5 and zebrafish KCNE6 but is not particularly similar to any single isoform (Fig. 1). Therefore, this subunit cannot be confidently assigned to a specific KCNE isoform based on sequence alone, and we use “KCNE0” to denote this lamprey KCNE subunit. Importantly, this naming does not imply a strict one-to-one evolutionary relationship with any particular jawed-vertebrate isoform.

Using deposited RNA-seq data from Far Eastern brook lamprey^27^ and RT-PCR in Arctic lamprey, we detected transcripts of both *kcnq1* and *kcne0* in multiple organs (Fig. 1E, F). Although transcript detection by RNA-seq or RT-PCR does not directly demonstrate protein expression, subcellular localization, or native complex assembly, the broad co-expression of transcripts supports the possibility that KCNQ1–KCNE0 complexes could operate in multiple lamprey tissues *in vivo*. This widespread expression contrasts with the more restricted and isoform-specific functions of KCNE subunits in humans, such as KCNE1, which is essential for ventricular repolarization and inner-ear K⁺ homeostasis, and KCNE3, which supports epithelial K⁺ recycling^1,2,7^. In this context, it is notable that *KCNE*-like genes can show lineage-specific gains, losses, and pseudogenization. For example, KCNE6 is functional in lower jawed vertebrates, including marsupials, but becomes a pseudogene in eutherians^10^, and a KCNE1-related pseudogene (KCNE1P) has been reported in zebra finch^44^. Consistent with these observations, one possible interpretation is that KCNE0 represents a relatively unspecialized KCNE subunit whose function and expression remain less restricted across tissues than those observed in jawed vertebrates, potentially allowing flexible pairing with KCNQ1 in different cellular contexts. Accordingly, in this study, we focus on the biophysical properties of KCNE0, particularly its effects on KCNQ1 in a heterologous expression system, while questions regarding its native physiological roles in lamprey tissues are left for future work.

When co-expressed with lamprey KCNQ1, KCNE0 produced a constitutively active current, resembling the effect typically induced by KCNE3 on mammalian KCNQ1^1,2,7^ (Fig. 2A–G; Supplementary Table 1). Our voltage-clamp fluorometry (VCF) experiments further support that KCNE0 modulates lamprey KCNQ1 gating by altering voltage-sensing domain (VSD) movement, consistent with how human KCNE subunits modulate KCNQ1 gating^31,33,34,36–39^ (Fig. 2H–J; Supplementary Table 1). An important observation is that KCNQ1–KCNE function is strongly dependent on the pairing of the two partners. In the native lamprey pairing, KCNE0 produced a constitutively active current accompanied by an apparent negative shift of the G–V relationship, whereas cross-species combinations showed weaker, mixed, or no effects on KCNQ1 modulation (Fig. 4; Supplementary Table 1). Previous cryo-electron microscopic structures^45–47^, together with a subsequent biophysical analysis^39^ showed extensive contacts between the KCNE transmembrane segment and the first transmembrane segment (S1) of KCNQ1 in the KCNQ1–KCNE complexes. However, the S1 segment is highly conserved among KCNQ1 orthologues across species. In contrast, the KCNQ1 proteins used in this study vary in amino-acid length, largely reflecting differences in the cytoplasmic C-terminal region (HsKCNQ1, 676 amino acids; DrKCNQ1, 575 amino acids; PmKCNQ1, 642 amino acids; LrKCNQ1, 644 amino acids; LcKCNQ1, 644 amino acids; CiKCNQ1, 520 amino acids) (Supplementary Fig. 4). Therefore, the species-specific effects observed in cross-species pairing experiments are unlikely to be explained by KCNE0 alone. Instead, they probably reflect compatibility between KCNE0 and species-specific features of KCNQ1 α-subunits, including regions beyond the conserved S1 segment, subtle differences within the broader transmembrane interface, and/or differences in intracellular regions that influence channel gating and coupling to KCNE subunits. Thus, sequence similarity around S1 alone is not sufficient to predict functional outcomes across distant lineages, and future structure-guided analyses will be needed to identify the KCNQ1-side determinants of KCNE compatibility.

Another mechanistic insight from this study is that KCNE0 can be shifted toward a KCNE4-like inhibitory effect by changes in a short intracellular region. Guided by prior work^41^ showing that KCNE4 contains a juxtamembrane tetra-leucine sequence required for interaction with calmodulin and functional suppression of KCNQ1 (Fig. 5A, B), introducing leucine substitutions that create a local tetra-leucine stretch in the corresponding region of KCNE0 markedly reduced KCNQ1 current amplitude and shifted the G–V relationship toward that of KCNQ1 expressed alone (Fig. 5C–F; Supplementary Table 1). One possible explanation for the ability of KCNE0 to acquire a KCNE4-like inhibitory effect is the presence of an extended intracellular region. Among KCNE family members, KCNE4 is distinguished by a long cytoplasmic tail that is essential for its inhibitory action on KCNQ1 (Fig. 1A), including interactions mediated by a juxtamembrane tetra-leucine motif^41^. KCNE0 similarly possesses a relatively long intracellular region compared with other KCNE isoforms (Fig. 1A). This shared architectural feature may influence KCNQ1-CaM interaction in a manner similar to KCNE4, thereby enabling KCNE0 to adopt KCNE4-like inhibitory properties. Taken together, these observations suggest that KCNE0 may represent a functionally flexible KCNE subunit that has not yet reached the degree of specialization observed among KCNE isoforms in jawed vertebrates, possibly reflecting an ancestral state preceding lineage-specific subfunctionalization of KCNE genes.

Several limitations of this study define clear next steps. First, we characterized KCNE0 primarily as an auxiliary subunit for KCNQ1 because KCNQ1 is the best-characterized pore-forming α-subunit partner for KCNE proteins. However, previous studies have shown that KCNE subunits can modulate other voltage-gated K^+^ channels^48–53^ and a Ca^2+^-gated Cl^-^ channel (TMEM16A)^54^ in heterologous expression systems, although at least one study failed to detect modulation of TMEM16A by KCNE1 under comparable conditions^55^. Second, our expression evidence is transcript-based and does not yet demonstrate protein expression, subcellular localization, or *in vivo* complex formation. Therefore, future studies will be required to test whether KCNE0 is functional *in vivo* and to identify its physiological partner(s) and contexts. Nevertheless, the broad tissue distribution of *kcne0* transcripts, together with the ability of KCNE0 to render KCNQ1 constitutively active, similar to the effect of mammalian KCNE3, raises the possibility that KCNQ1–KCNE0 complexes contribute to general ion homeostasis, possibly including K^+^ recycling in epithelial tissues, rather than to a highly specialized role like that of KCNQ1–KCNE1 in the mammalian heart and inner ear. Third, while cross-species pairing experiments support species-specific tuning of compatibility, we do not yet know which precise interface features encode compatibility across lineages. Addressing these questions will require additional structure-guided biophysical analyses.

In summary, these findings provide a framework for comparative studies of KCNE-dependent KCNQ1 modulation across vertebrate lineages and suggest that KCNE0 represents a relatively unspecialized KCNE subunit, potentially reflecting an ancestral stage preceding lineage-specific subfunctionalization of KCNE genes in jawed vertebrates.

## Acknowledgements

We thank the members of the Nakajo laboratory for helpful discussions and support. This work was supported by the Japan Society for the Promotion of Science KAKENHI (Grant Nos. 23K27357 to G.K., 24K09531 to E.K.Y., and 24K02211 to K.N.).

## Author contributions

G.K. and K.N. conceived and designed the project. G.K. and K.R. performed electrophysiological experiments. G.K. performed confocal microscopy. G.K., B.Z., E.K.Y., and K.N. collected tissue samples and performed RT-PCR. G.K. analyzed deposited RNA-seq data. G.K. and K.N. wrote the original draft, and all authors reviewed and edited the manuscript. G.K. and K.N. supervised the research.

## Competing interests

The authors declare no competing interests.

## Data availability

All data supporting this study are available within the paper and its Supplementary Information. Additional materials are available from the corresponding author upon reasonable request.

## Materials and Methods

### Protein expression in *Xenopus laevis* oocytes

The coding regions of KCNQ1 (human, NCBI Accession Number NM_000218.3; zebrafish, NM_001123242.2; sea lamprey, XM_076065335.1; Far Eastern brook lamprey, XM_061564454.1; Arctic lamprey, see Supplementary Fig. 1; vase tunicate, NM_001160065), KCNE0 (sea lamprey, XM_032975225.1; Far Eastern brook lamprey, XM_061575561.1; Arctic lamprey, see Supplementary Fig. 2), human KCNE1 (NM_000219.6), and human KCNE3 (NM_005472.5) were cloned into the pGEMHE vector^56^. cRNA was transcribed using the HiScribe^®^ T7 ARCA mRNA Kit (New England Biolabs, E2065S). Female *Xenopus laevis* frogs were anesthetized in water containing 0.1% tricaine (Sigma-Aldrich, E10521) for 15–30 min, and oocytes were surgically isolated. Follicle layers were removed by collagenase treatment (Sigma-Aldrich, C0130) for 5–6 h at room temperature. Defolliculated oocytes of similar size at stage V–VI were selected, microinjected with 50 nL of cRNA solution (2–10 ng for KCNQ1 and 1 ng for KCNE) using a Nanoject II (Drummond Scientific), and incubated at 18 °C in Barth’s solution (88 mM NaCl, 1 mM KCl, 2.4 mM NaHCO_3_, 10 mM HEPES, 0.3 mM Ca(NO_3_)_2_, 0.41 mM CaCl_2_, and 0.82 mM MgSO_4_, pH 7.6) supplemented with 0.1% penicillin -streptomycin solution (Sigma-Aldrich, P4333). All procedures involving *Xenopus laevis* were approved by the Animal Care Committee of Jichi Medical University (protocol 21030-04) and complied with institutional guidelines.

### Two-electrode voltage clamp (TEVC) recordings

Oocytes were recorded 1–3 days after injection using an OC-725C amplifier (Warner Instruments) at room temperature. The bath was continuously perfused with Ca^2+^-free ND96 (96 mM NaCl, 2 mM KCl, 2.8 mM MgCl_2_, 5 mM HEPES, pH 7.6) containing 100 µM LaCl_3_ to suppress endogenous hyperpolarization-activated currents^29,35,39^. Microelectrodes were pulled from borosilicate glass capillaries (Harvard Apparatus, GC150TF-10) using a P-1000 micropipette puller (Sutter Instrument) to 0.2–1.0 MΩ and filled with 3 M KCl. From a holding potential of -90 mV, currents were elicited by voltage steps from −100 to +60 mV in +20 mV increments with 2 s step duration and 10 s intervals. Oocytes with a holding current < -0.4 µA at -90 mV were excluded. Protocol generation and data acquisition were performed using a Digidata 1550 (Molecular Devices) controlled by pCLAMP 10.7. Signals were sampled at 10 kHz and low-pass filtered at 1 kHz.

### Voltage dependence analysis (G-V)

G–V relationships were obtained from tail current amplitudes at -30 mV. Fits were performed in pCLAMP 10.7 (Molecular Devices) to a single-Boltzmann function:

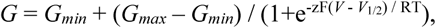

where *G_max_* and *G_min_* are the maximal and minimal tail conductances, *z* is the effective gating charge, *V_1/2_* is the half-activation voltage, T is absolute temperature, F is Faraday’s constant, and R is the gas constant. Normalized conductance (*G*/*G_max_*) was plotted against voltage for presentation.

### Voltage-clamp fluorometry

Oocytes expressing constructs for VCF were incubated for 2–4 days after injection. Labeling was performed for 30 min in KD98 solution (98 mM KCl, 1.8 mM CaCl_2_, 1 mM MgCl_2_, 5 mM HEPES, pH 7.6) with 5 µM Alexa Fluor^TM^ 488 C_5_ maleimide (Thermo Fisher Scientific, A10254), and unreacted dye was removed by washing with Ca^2+^-free ND96 solution^29,35,39^. Microelectrodes were pulled from borosilicate glass capillaries (Harvard Apparatus, GC150TF-15), as in TEVC recordings. From a holding potential of -90 mV, currents and fluorescence signals were recorded during voltage steps from +80 to -180 mV, or from +60 mV to -140 mV where indicated, in -20 mV increments with 2 s step duration and 20 s intervals. Oocytes with a holding current < -0.4 µA at -90 mV were excluded, as in TEVC recordings. Protocol generation and data acquisition were performed using a Digidata 1440A (Molecular Devices) controlled by pCLAMP 10.7. Ionic currents were sampled at 10 kHz and low-pass filtered at 1 kHz. Fluorescence signals were digitized at 1 kHz through the Digidata1440A and low-pass filtered at 50 Hz.

Fluorescence recordings were obtained with an MVX10 macrozoom microscope (Olympus) equipped with a 2x objective lens (MVPLAPO 2XC, NA = 0.5, Olympus), a 2x magnification changer (MVX-CA2X, Olympus), a GFP filter cube (U-MGFPHQ/XL, Olympus), and an XLED1 light source with a BDX 450-495 nm LED module (Excelitas Technologies). Fluorescence was detected with a photomultiplier (H10722-110, Hamamatsu Photonics) and recorded in pCLAMP 10.7 simultaneously with ionic currents. The excitation shutter remained open during recordings, which caused a gradual decrease in fluorescence due to photobleaching. For each trace, the bleaching rate (R) was estimated from the 1.1 s baseline preceding the test pulse and traces were corrected assuming a linear decrease:

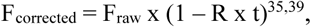

Corrected traces were then baseline-normalized to 1 at the pre-step level.

### VCF analysis

F–V relationships were obtained by plotting the fluorescence change from the baseline (ΔF) against membrane potential. For presentation (Fig. 2), ΔF values were normalized to the response at +80 mV (ΔF_+80mV_). Fits were performed in Igor Pro (WaveMetrics). KCNQ1 alone was fit with a double-Boltzmann function, whereas KCNQ1 co-expressed with KCNE0 was fit with a single-Boltzmann function.

Single-Boltzmann function:

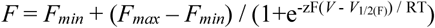

Double-Boltzmann function:

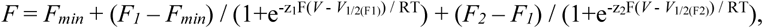

where *F_min_*, *F_1_*, *F_2_*, and *F_max_* denote the baseline, intermediate, and maximal fluorescence components, zF is the effective gating charge for the fluorescence component, *V_1/2_*_(F)_, *V_1/2_*_(F1)_, and *V_1/2_*_(F2)_ are the half-activation voltages for the fluorescence components, T is absolute temperature, F is Faraday’s constant, and R is the gas constant. Normalized fluorescence change (*ΔF*/*ΔF_+80mV_*) was plotted against voltage for presentation.

### Isolation of cDNAs, cloning, and RT-PCR from Arctic lamprey

Arctic lampreys (*Lethenteron camtschaticum*) captured in the Ishikari River (Hokkaido, Japan) were commercially obtained and maintained on 12-h light/12-h dark cycles at 4 °C. Total RNA was extracted from the indicated organs using NucleoSpin RNA Plus (MACHEREY-NAGEL, 740984) and reverse-transcribed using PrimeScript II (TaKaRa, 6210) according to the manufacturer’s instructions. The resulting cDNA was used as a template for PCR amplification. PCR for cloning was performed using KOD One PCR Master Mix (TOYOBO, KMM-101), and RT-PCR of β-actin, KCNQ1, and KCNE0 was performed using PrimeSTAR Max DNA Polymerase (TaKaRa, R045A), according to the manufacturers’ protocols.

For cloning of KCNQ1 and KCNE0, we performed two-step PCR using the following primers

KCNQ1 1st Fw 5′-ATGTCACACGGAAAGCGAAGTTCTTCTCACAGAGG-3′

KCNQ1 1st Rv 5′-ACAGCTGTGCTGTTGGCTGAAAGGTATGTGGGCGC-3′

KCNQ1 2nd Fw 5′-AGTGGCGGAGCCACCATGTCACACGGAAAGCGAAG-3′

KCNQ1 2nd Rv 5′-GTCGCGGCCGCTTTAACAGCTGTGCTGTTGGCTGA-3′

KCNE0 1st Fw 5′-GACACGGAGAGAGCGAGCGCCGGCGAC-3′

KCNE0 1st Rv 5′-AGGGGCTGGAGGTTAGGAGCTGGGCCC-3′

KCNE0 2nd Fw 5′-AGTGGCGGAGCCACCGACACGGAGAGAGCGAGCGC-3′

KCNE0 2nd Rv 5′-GTCGCGGCCGCTTTAAGGGGCTGGAGGTTAGGAGC-3′

For RT-PCR we used the following primers

β-actin Fw 5′-ACCCAGATCATGTTTGAGACC-3′

β-actin Rv 5′-GACTCCATGCCGATGAATGA-3′

KCNQ1 Fw 5′-CCTGGGTCTCATATTCTCATC-3′

KCNQ1 Rv 5′-TGACATCTCCACAGGCTCTG-3′

KCNE0 Fw 5′-ACATGCAGGGCCTCTCATCG-3′

KCNE0 Rv 5′-TGCACGTAGAGGTGGAACGG-3′

### Analysis of deposited RNA-seq data from Far Eastern brook lamprey

The raw paired-end RNA-seq FASTQ files were downloaded from the Sequence Read Archive (SRA). The run identifiers were SRR9964076 (heart), SRR9964077 (gill), SRR9964078 (testis), SRR9964079 (brain), SRR9964080 (liver), SRR9964081 (oral gland), SRR9964082 (kidney), SRR9964083 (intestine), SRR9964084 (supraneural body), and SRR9964085 (muscle). Raw reads were processed for adapter removal, base-quality filtering, and per-read quality evaluation using fastp (v1.1.0)^57,58^. Quality control reports were generated with FastQC (v0.12.1) and MultiQC (v1.33)^59^ (Supplementary Table 2). A decoy-aware Salmon index was built from two target transcripts (KCNQ1, XM_061564454.1; KCNE0, XM_061575561.1) combined with RefSeq decoy sequences (GCF_015708825.1). Quantification was performed with Salmon (v1.10.3)^60^ (Supplementary Table 3). Transcript-level estimates were summarized at the gene level and reported as counts per million (CPM).

### Statistics and reproducibility

Data are presented as mean ± SEM (n = 5–8). In Figs. 2G, 3F, and 3N, differences in G–V *V_1/2_* values among groups were assessed using one-way ANOVA followed by the Tukey–Kramer multiple-comparison test to evaluate pairwise differences. In Fig. 5F, differences in current amplitudes among groups were assessed using one-way ANOVA followed by the Tukey–Kramer multiple-comparison test to evaluate pairwise differences. In Figs. 4C, 4F, 4I, and 4L, differences between two groups were assessed using unpaired two-tailed Welch’s t-test. *V_1/2_* values were not statistically compared when reliable Boltzmann fits could not be obtained because the G–V relationships were nearly saturated over the tested voltage range. Statistical significance was defined as P < 0.05 (*P < 0.05, **P < 0.01, *** P < 0.001).

---

**Supplementary Figure 1.**
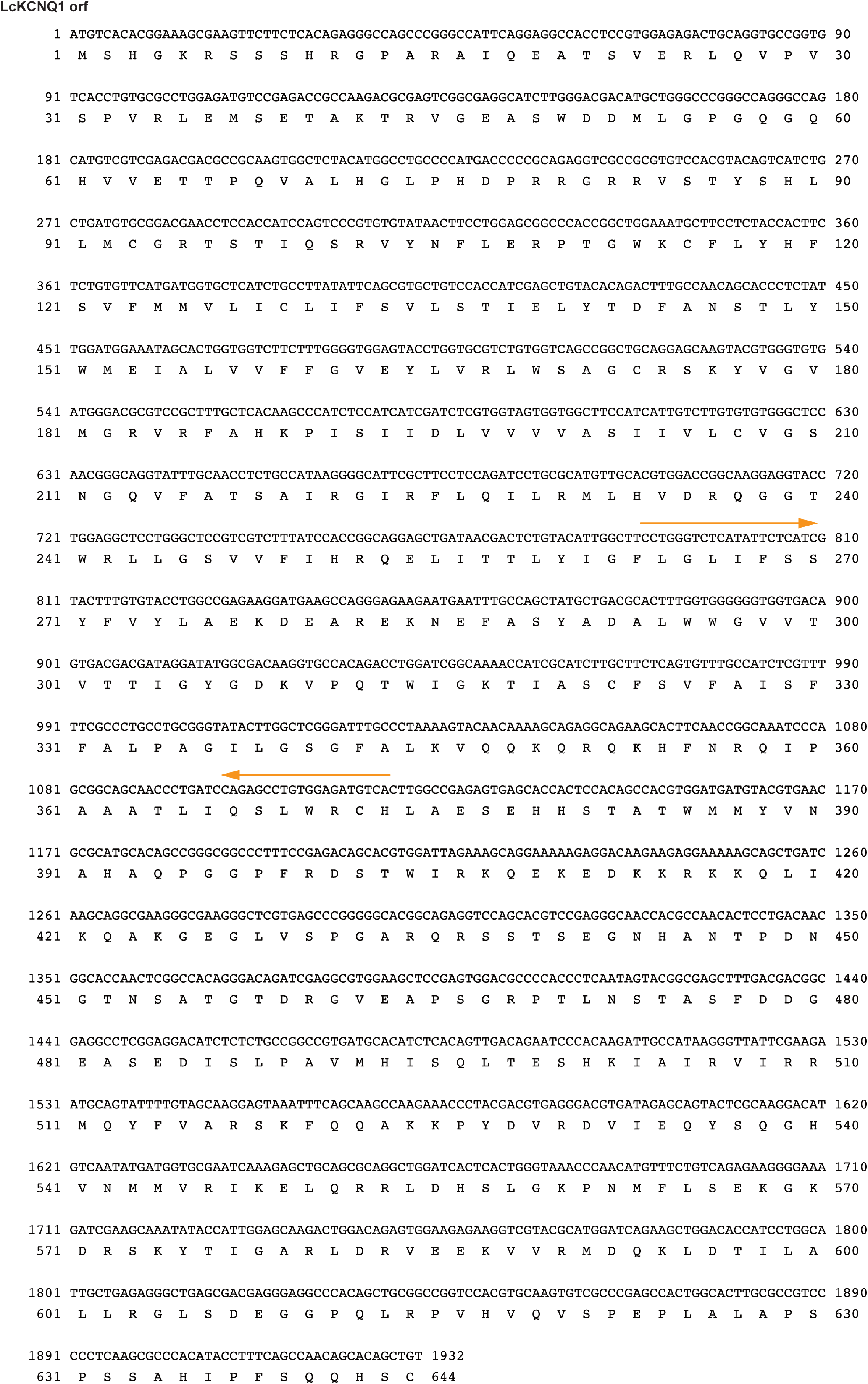
ORF nucleotide and corresponding amino-acid sequence of KCNQ1 in Arctic lamprey. The open reading frame (ORF) nucleotide sequence and the translated amino-acid sequence of Arctic lamprey KCNQ1 used for cloning and electrophysiological recordings are shown. The primer-binding sites used for RT-PCR analysis in Fig. 1F are indicated with orange arrows.

**Supplementary Figure 2.**
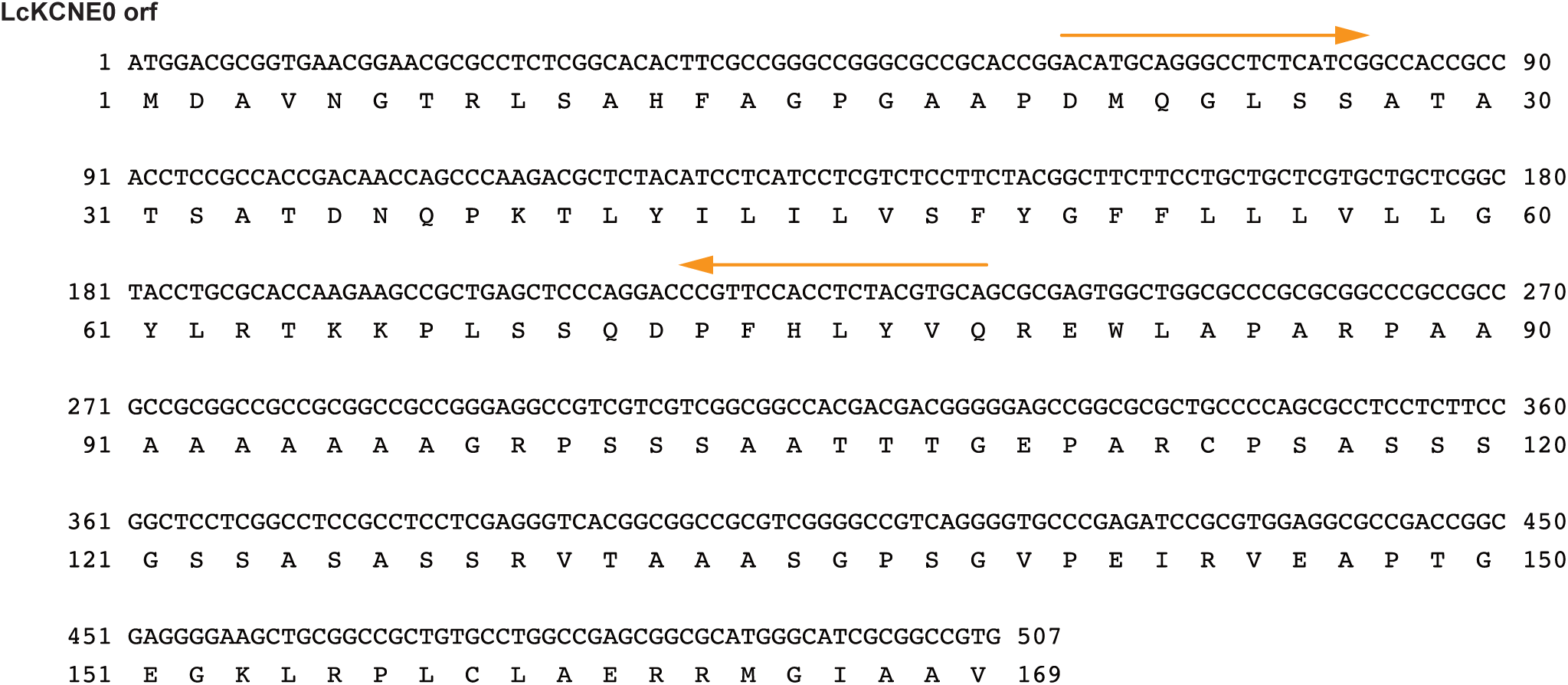
ORF nucleotide and corresponding amino-acid sequence of KCNE0 in Arctic lamprey. The open reading frame (ORF) nucleotide sequence and the translated amino-acid sequence of Arctic lamprey KCNE0 used for cloning and electrophysiological recordings are shown. The primer-binding sites used for RT-PCR analysis in Fig. 1F are indicated with orange arrows.

**Supplementary Figure 3.**
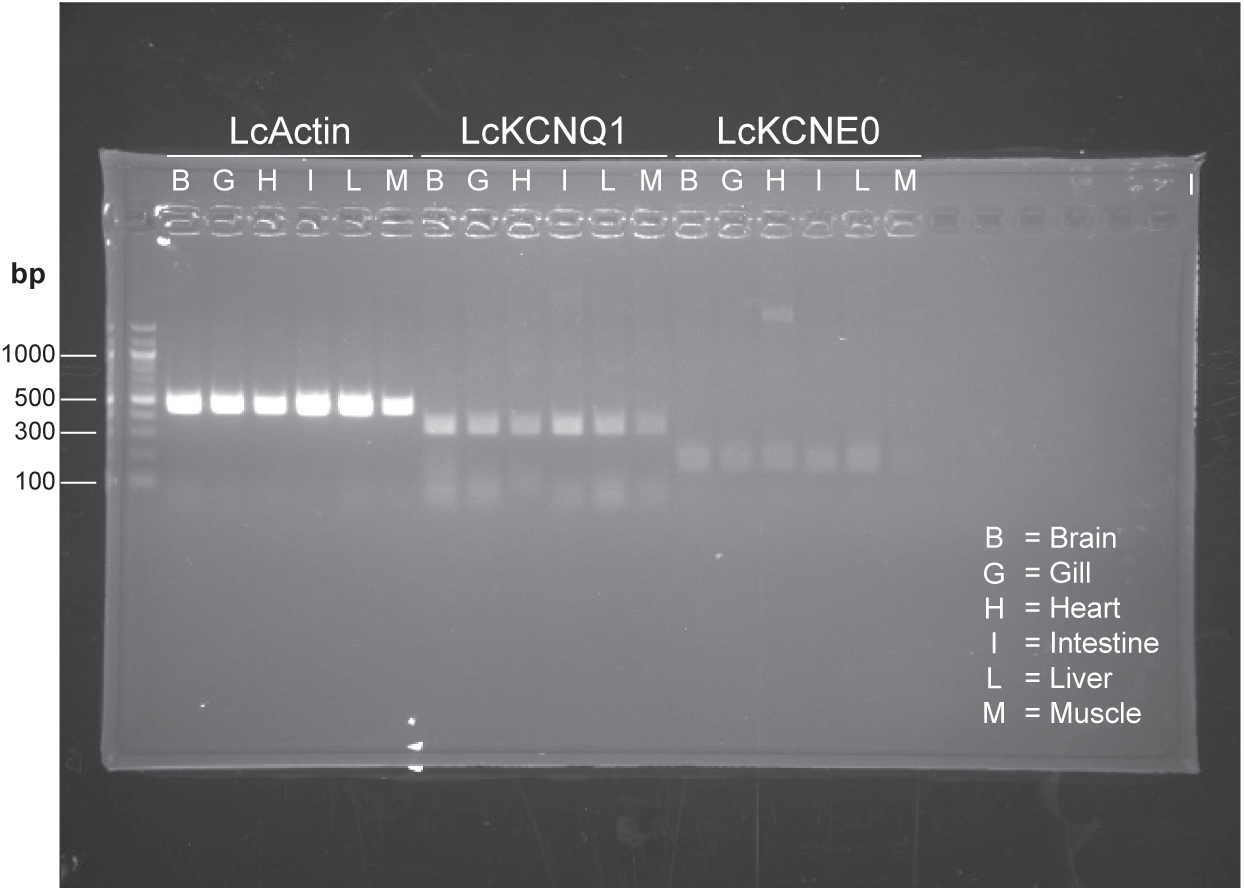
Uncropped agarose gel corresponding to the gel image shown in Figure 1F.

**Supplementary Figure 4.**
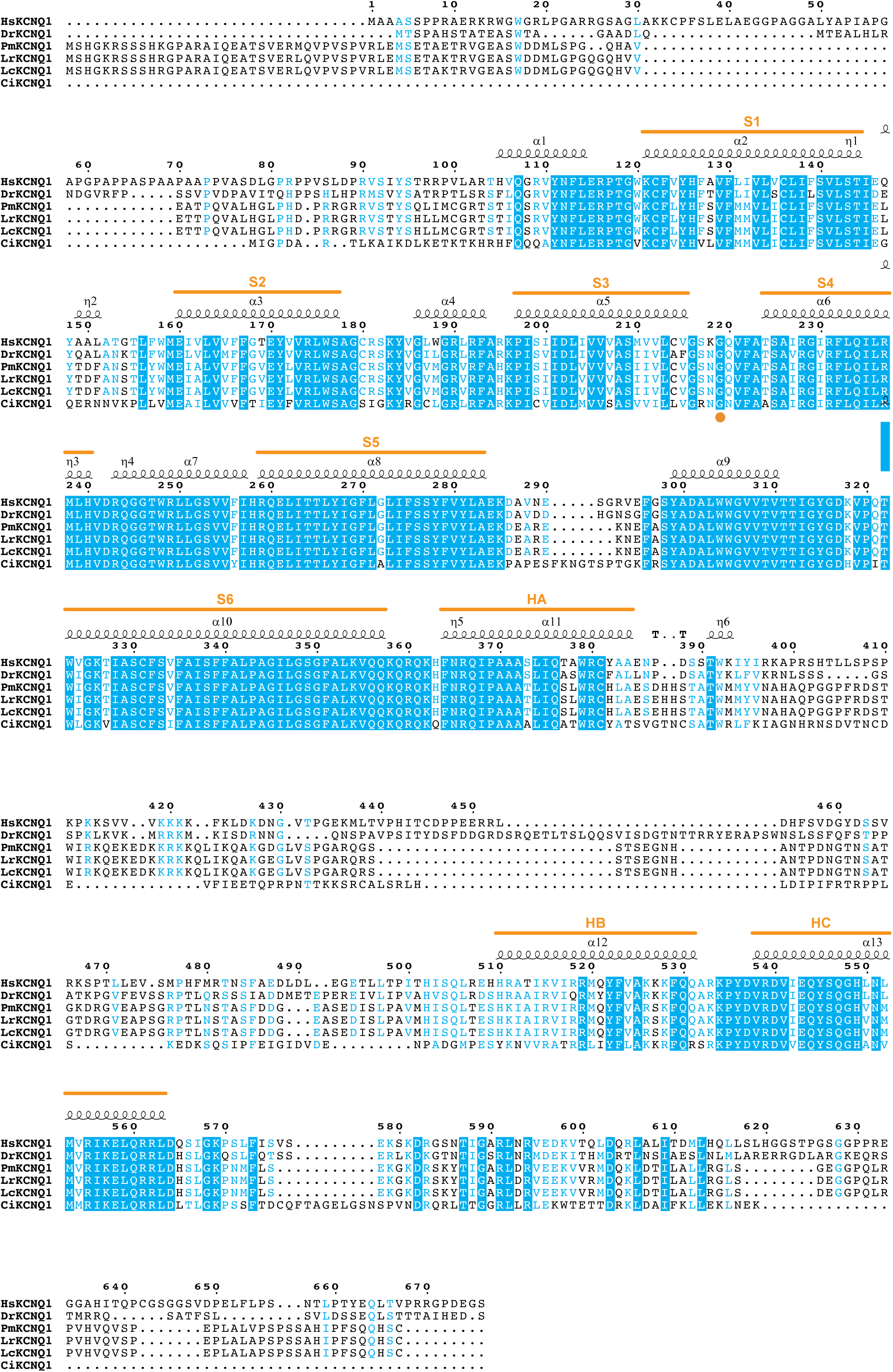
Amino-acid sequence alignment of KCNQ1 from various species. Amino acid sequence alignment of KCNQ1 from human, zebrafish, lampreys, and vase tunicate generated with Clustal Omega^61^ and displayed with ESPript3^62^. The residue corresponding to the labeling site in human KCNQ1 (G219)^29,30^ is highlighted with an orange circle. For sequence alignment, human (HsKCNQ1; NCBI Accession Number NP_000209.2), zebrafish (DrKCNQ1; NP_001116714.1), sea lamprey (PmKCNQ1; XP_075921450.1), Far Eastern brook lamprey (LrKCNQ1; XP_061420438.1), Arctic lamprey, (LcKCNQ1; see Supplementary Fig. 1), and vase tunicate (CiKCNQ1; NP_001153537.1) KCNQ1 were used.

**Supplementary Figure 5.**
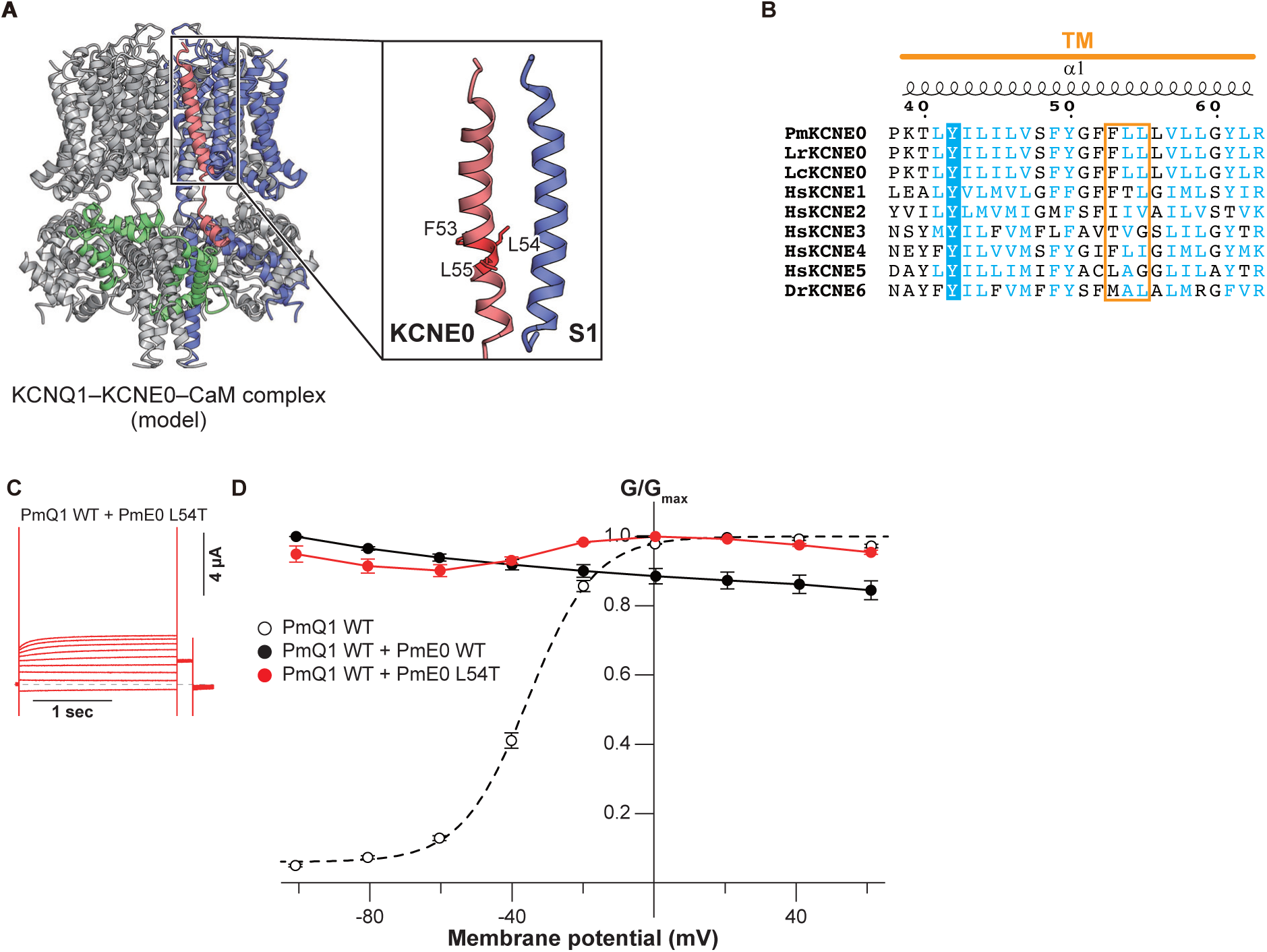
Attempted conversion of KCNE0 toward a KCNE1-like effect. (**A**) Close-up view of the interface between KCNQ1 and KCNE0 within the PmKCNQ1–PmKCNE0–PmCaM complex corresponding to the structure shown in Fig. 5A. (**B**) Sequence alignment around the KCNE transmembrane segments highlighting the “triplet” region^42,43^. (**C,D**) A representative current trace (C) and G-V relationship (D) of PmKCNQ1 WT co-expressed with the HsKCNE1-like triplet mutant of PmKCNE0 (PmKCNE0 L54T). In (D), error bars denote mean ± s.e.m. for n = 5.

**Supplementary Table 1. Summary of electrophysiological parameters and V1/2 statistical comparisons.** Summary of electrophysiological parameters for all constructs. For each condition, maximum tail current amplitude (*I_max_*) and Boltzmann fit parameters (*V_1/2_* and *z*) are reported. For VCF experiments, parameters from single- or double-Boltzmann fits are reported as appropriate. Values are mean ± s.e.m.; n indicates the number of oocytes. In the statistical comparisons, P values indicate pairwise comparisons of G–V *V_1/2_* values. For datasets containing more than two groups (Figs. 2G, 3F, and 3N), one-way ANOVA was first performed, followed by Tukey–Kramer multiple-comparison tests. For two-group comparisons (Fig. 4C, 4F, 4I, and 4L), an unpaired two-tailed Welch’s t-test was used. *V_1/2_* values were not statistically compared when reliable Boltzmann fits could not be obtained because the G–V relationships were nearly saturated over the tested voltage range.

**Supplementary Table 2. Summary of fastp quality control.** Quality-control statistics generated by fastp for all RNA-seq runs. For each run, total reads before and after trimming (million reads), percentage of reads retained after trimming (%), mean read length of R1 and R2 before and after trimming (bp), and the percentage of bases with Phred quality ≥ 30 (Q30) before and after trimming (%) are shown. “Reads” denote paired-end fragments.

**Supplementary Table 3. Quantification summary using Salmon.** Summary of transcript detections for Far Eastern brook lamprey KCNQ1 (XM_061564454.1) and KCNE0 (XM_061575561.1) using Salmon. For each run, the number of detected fragments, total processed fragments, and CPM (counts per million of the library) are shown. “Fragments” denote paired-end read pairs (one read pair counted as one fragment).

**Source data. Excel file with numerical electrophysiology data acquired in this study.**

## Notes

### Competing Interest Statement

The authors have declared no competing interest.

### Summary of Updates

Addition of Statistical Analyses and Enhanced Data Visualization: In accordance with the reviewers' recommendations, we have performed additional statistical analyses to compare the V1/2 values. Furthermore, the error bars in the figures have been modified to improve visual clarity and readability. Expansion of the Discussion on Physiological Roles: We have expanded and revised the Discussion section to include further insights and speculations regarding the physiological roles of our findings, directly addressing the points raised during the review.

## References

1. Abbott, G. W. KCNE1 and KCNE3: The yin and yang of voltage-gated K(+) channel regulation. Gene 576, 1–13 (2016).

2. Abbott, G. W. Kv Channel Ancillary Subunits: Where Do We Go from Here? Physiology 37, 225–241 (2022).

3. Takumi, T., Ohkubo, H. & Nakanishi, S. Cloning of a membrane protein that induces a slow voltage-gated potassium current. Science 242, 1042–1045 (1988).

4. Abbott, G. W. et al. MiRP1 forms IKr potassium channels with HERG and is associated with cardiac arrhythmia. Cell 97, 175–187 (1999).

5. Piccini, M. et al. KCNE1-like gene is deleted in AMME contiguous gene syndrome: identification and characterization of the human and mouse homologs. Genomics 60, 251–257 (1999).

6. Tinel, N., Diochot, S., Borsotto, M., Lazdunski, M. & Barhanin, J. KCNE2 confers background current characteristics to the cardiac KCNQ1 potassium channel. EMBO J. 19, 6326–6330 (2000).

7. Schroeder, B. C. et al. A constitutively open potassium channel formed by KCNQ1 and KCNE3. Nature 403, 196–199 (2000).

8. Grunnet, M. et al. KCNE4 is an inhibitory subunit to the KCNQ1 channel. J. Physiol. 542, 119–130 (2002).

9. Angelo, K. et al. KCNE5 induces time- and voltage-dependent modulation of the KCNQ1 current. Biophys. J. 83, 1997–2006 (2002).

10. Kasuya, G. et al. Identification of KCNE6, a new member of the KCNE family of potassium channel auxiliary subunits. *Commun*. Biol. 7, 1662 (2024).

11. Barhanin, J. et al. KvLQT1 and lsK (minK) proteins associate to form the IKs cardiac potassium current. Nature 384, 78–80 (1996).

12. Sanguinetti, M. C. et al. Coassembly of K(V)LQT1 and minK (IsK) proteins to form cardiac I(Ks) potassium channel. Nature 384, 80–83 (1996).

13. Neyroud, N. et al. A novel mutation in the potassium channel gene KVLQT1 causes the Jervell and Lange-Nielsen cardioauditory syndrome. Nat. Genet. 15, 186–189 (1997).

14. Tyson, J. et al. IsK and KvLQT1: mutation in either of the two subunits of the slow component of the delayed rectifier potassium channel can cause Jervell and Lange-Nielsen syndrome. Hum. Mol. Genet. 6, 2179–2185 (1997).

15. Abbott, G. W. The KCNE2 K+ channel regulatory subunit: Ubiquitous influence, complex pathobiology. Gene 569, 162–172 (2015).

16. Abbott, G. W. KCNE4 and KCNE5: K(+) channel regulation and cardiac arrhythmogenesis. Gene 593, 249–260 (2016).

17. Vallon, V. et al. Role of KCNE1-dependent K+ fluxes in mouse proximal tubule. J. Am. Soc. Nephrol. 12, 2003–2011 (2001).

18. Finley, M. R. et al. Expression and coassociation of ERG1, KCNQ1, and KCNE1 potassium channel proteins in horse heart. Am. J. Physiol. Heart Circ. Physiol. 283, H126–38 (2002).

19. Morokuma, J., Blackiston, D. & Levin, M. KCNQ1 and KCNE1 K+ channel components are involved in early left-right patterning in Xenopus laevis embryos. Cell. Physiol. Biochem. 21, 357–372 (2008).

20. Abramochkin, D. V., Hassinen, M. & Vornanen, M. Transcripts of Kv7.1 and MinK channels and slow delayed rectifier K+ current (IKs) are expressed in zebrafish (Danio rerio) heart. Pflugers Arch. 470, 1753–1764 (2018).

21. Haverinen, J., Hassinen, M. & Vornanen, M. Effect of Channel Assembly (KCNQ1 or KCNQ1 + KCNE1) on the Response of Zebrafish IKs Current to IKs Inhibitors and Activators. J. Cardiovasc. Pharmacol. 79, 670–677 (2022).

22. Okamura, Y. et al. Comprehensive analysis of the ascidian genome reveals novel insights into the molecular evolution of ion channel genes. Physiol. Genomics 22, 269–282 (2005).

23. Park, K. H., Hernandez, L., Cai, S.-Q., Wang, Y. & Sesti, F. A family of K+ channel ancillary subunits regulate taste sensitivity in Caenorhabditis elegans. J. Biol. Chem. 280, 21893–21899 (2005).

24. Fenyves, B. G. et al. Dual role of an mps-2/KCNE-dependent pathway in long-term memory and age-dependent memory decline. Curr. Biol. 31, 527–539.e7 (2021).

25. Heimberg, A. M., Cowper-Sal-lari, R., Sémon, M., Donoghue, P. C. J. & Peterson, K. J. microRNAs reveal the interrelationships of hagfish, lampreys, and gnathostomes and the nature of the ancestral vertebrate. Proc. Natl. Acad. Sci. U. S. A. 107, 19379–19383 (2010).

26. Stock, D. W. & Whitt, G. S. Evidence from 18S ribosomal RNA sequences that lampreys and hagfishes form a natural group. Science 257, 787–789 (1992).

27. Zhu, T. et al. Chromosome-level genome assembly of Lethenteron reissneri provides insights into lamprey evolution. Mol. Ecol. Resour. 21, 448–463 (2021).

28. Cowgill, J. & Chanda, B. The contribution of voltage clamp fluorometry to the understanding of channel and transporter mechanisms. J. Gen. Physiol. jgp.201912372 (2019).

29. Osteen, J. D. et al. KCNE1 alters the voltage sensor movements necessary to open the KCNQ1 channel gate. Proc. Natl. Acad. Sci. U. S. A. 107, 22710–22715 (2010).

30. Osteen, J. D. et al. Allosteric gating mechanism underlies the flexible gating of KCNQ1 potassium channels. Proc. Natl. Acad. Sci. U. S. A. 109, 7103–7108 (2012).

31. Zaydman, M. A. et al. Domain–domain interactions determine the gating, permeation, pharmacology, and subunit modulation of the IKs ion channel. Elife 3, e03606 (2014).

32. Hou, P. et al. Inactivation of KCNQ1 potassium channels reveals dynamic coupling between voltage sensing and pore opening. Nat. Commun. 8, 1–11 (2017).

33. Hou, P. et al. Two-stage electro-mechanical coupling of a KV channel in voltage-dependent activation. Nat. Commun. 11, 676 (2020).

34. Barro-Soria, R. et al. KCNE1 divides the voltage sensor movement in KCNQ1/KCNE1 channels into two steps. Nat. Commun. 5, 3750 (2014).

35. Nakajo, K. & Kubo, Y. Steric hindrance between S4 and S5 of the KCNQ1/KCNE1 channel hampers pore opening. Nat. Commun. 5, 4100 (2014).

36. Barro-Soria, R., Perez, M. E. & Larsson, H. P. KCNE3 acts by promoting voltage sensor activation in KCNQ1. Proc. Natl. Acad. Sci. U. S. A. 112, E7286–E7292 (2015).

37. Barro-Soria, R. et al. KCNE1 and KCNE3 modulate KCNQ1 channels by affecting different gating transitions. Proc. Natl. Acad. Sci. U. S. A. 114, E7367–E7376 (2017).

38. Taylor, K. C. et al. Structure and physiological function of the human KCNQ1 channel voltage sensor intermediate state. Elife 9, 1–31 (2020).

39. Kasuya, G. & Nakajo, K. Optimized tight binding between the S1 segment and KCNE3 is required for the constitutively open nature of the KCNQ1-KCNE3 channel complex. Elife 11, 1–19 (2022).

40. Nakajo, K., Nishino, A., Okamura, Y. & Kubo, Y. KCNQ1 subdomains involved in KCNE modulation revealed by an invertebrate KCNQ1 orthologue. J. Gen. Physiol. 138, 521–535 (2011).

41. Ciampa, E. J., Welch, R. C., Vanoye, C. G. & George, A. L., Jr. KCNE4 juxtamembrane region is required for interaction with calmodulin and for functional suppression of KCNQ1. J. Biol. Chem. 286, 4141–4149 (2011).

42. Melman, Y. F., Domènech, A., De la Luna, S. & McDonald, T. V. Structural Determinants of KvLQT1 Control by the KCNE Family of Proteins. J. Biol. Chem. 276, 6439–6444 (2001).

43. Melman, Y. F., Krumerman, A. & McDonald, T. V. A single transmembrane site in the KCNE-encoded proteins controls the specificity of KvLQT1 channel gating. J. Biol. Chem. 277, 25187–25194 (2002).

44. Lovell, P. V., Carleton, J. B. & Mello, C. V. Genomics analysis of potassium channel genes in songbirds reveals molecular specializations of brain circuits for the maintenance and production of learned vocalizations. BMC Genomics 14, 470 (2013).

45. Cui, C. et al. Mechanisms of KCNQ1 gating modulation by KCNE1/3 for cell-specific function. Cell Res. 35, 876–886 (2025).

46. Zhong, L. et al. Secondary structure transitions and dual PIP2 binding define cardiac KCNQ1-KCNE1 channel gating. Cell Res. 35, 887–899 (2025).

47. Sun, J. & MacKinnon, R. Structural Basis of Human KCNQ1 Modulation and Gating. Cell 180, 340–347 (2020).

48. McDonald, T. V. et al. A minK-HERG complex regulates the cardiac potassium current I(Kr). Nature 388, 289–292 (1997).

49. McCrossan, Z. A. et al. MinK-related peptide 2 modulates Kv2.1 and Kv3.1 potassium channels in mammalian brain. J. Neurosci. 23, 8077–8091 (2003).

50. Abbott, G. W. et al. MiRP2 forms potassium channels in skeletal muscle with Kv3.4 and is associated with periodic paralysis. Cell 104, 217–231 (2001).

51. Deschênes, I. & Tomaselli, G. F. Modulation of Kv4.3 current by accessory subunits. FEBS Lett. 528, 183–188 (2002).

52. Delpón, E. et al. Functional effects of KCNE3 mutation and its role in the development of Brugada syndrome. Circ. Arrhythm. Electrophysiol. 1, 209–218 (2008).

53. Wang, W. et al. Functional significance of K+ channel β-subunit KCNE3 in auditory neurons. J. Biol. Chem. 289, 16802–16813 (2014).

54. Ávalos Prado, P., et al. KCNE1 is an auxiliary subunit of two distinct ion channel superfamilies. Cell 184, 534–544.e11 (2021).

55. Talbi, K., Ousingsawat, J., Centeio, R., Schreiber, R. & Kunzelmann, K. KCNE1 does not shift TMEM16A from a Ca2+ dependent to a voltage dependent Cl- channel and is not expressed in renal proximal tubule. Pflugers Arch. 475, 995–1007 (2023).

56. Liman, E. R., Tytgat, J. & Hess, P. Subunit stoichiometry of a mammalian K+ channel determined by construction of multimeric cDNAs. Neuron 9, 861–871 (1992).

57. Chen, S., Zhou, Y., Chen, Y. & Gu, J. fastp: an ultra-fast all-in-one FASTQ preprocessor. Bioinformatics 34, i884–i890 (2018).

58. Chen, S. fastp 1.0: An ultra-fast all-round tool for FASTQ data quality control and preprocessing. Imeta 4, e70078 (2025).

59. Ewels, P., Magnusson, M., Lundin, S. & Käller, M. MultiQC: summarize analysis results for multiple tools and samples in a single report. Bioinformatics 32, 3047–3048 (2016).

60. Patro, R., Duggal, G., Love, M. I., Irizarry, R. A. & Kingsford, C. Salmon provides fast and bias-aware quantification of transcript expression. Nat. Methods 14, 417–419 (2017).

61. Madeira, F. et al. The EMBL-EBI Job Dispatcher sequence analysis tools framework in 2024. Nucleic Acids Res. 52, W521–W525 (2024).

62. Robert, X. & Gouet, P. Deciphering key features in protein structures with the new ENDscript server. Nucleic Acids Res. 42, 320–324 (2014).

63. Martin, F. J. et al. Ensembl 2023. Nucleic Acids Res. 51, D933–D941 (2023).

64. Waterhouse, A. et al. SWISS-MODEL: homology modelling of protein structures and complexes. Nucleic Acids Res. 46, W296–W303 (2018).

