## Supplemental Table 1 for "Characterization of an early-diverging KCNE potassium-channel auxiliary subunit in the jawless vertebrate lamprey"

| Conditions (TEVC) | $I_{max}$ (µA) | $V_{1/2}$ (mV) | *z* | *n* |
|---|---|---|---|---|
| **Figure 2** | | | | |
| PmKCNQ1 WT | 1.4 ± 0.2 | -35.8 ± 0.9 | 2.8 ± 0.1 | 5 |
| PmKCNQ1 WT + PmKCNE0 WT | *2.0 ± 0.1 | n.d. | n.d. | 5 |
| LrKCNQ1 WT | 1.6 ± 0.2 | -32.9 ± 0.6 | 2.8 ± 0.1 | 5 |
| LrKCNQ1 WT + LrKCNE0 WT | *2.1 ± 0.2 | n.d. | n.d. | 5 |
| LcKCNQ1 WT | 1.8 ± 0.2 | -32.1 ± 0.6 | 2.5 ± 0.1 | 5 |
| LcKCNQ1 WT + LcKCNE0 WT | *2.8 ± 0.3 | n.d. | n.d. | 5 |
| **Figure 3** | | | | |
| PmKCNQ1 WT–PmKCNE0ΔN6 | 1.7 ± 0.1 | -37.8 ± 0.7 | 2.9 ± 0.0 | 5 |
| PmKCNQ1 WT–PmKCNE0ΔN11 | 1.4 ± 0.2 | -37.9 ± 1.0 | 2.7 ± 0.1 | 5 |
| PmKCNQ1 WT–PmKCNE0ΔN22 | 1.6 ± 0.2 | -39.4 ± 0.8 | 3.2 ± 0.0 | 5 |
| PmKCNQ1 WT–PmKCNE0ΔN30 | 1.6 ± 0.1 | -34.3 ± 0.5 | 2.8 ± 0.1 | 5 |
| PmKCNQ1 WT–PmKCNE0ΔC30 | *2.5 ± 0.1 | n.d. | n.d. | 5 |
| PmKCNQ1 WT–PmKCNE0ΔC61 | *2.0 ± 0.4 | n.d. | n.d. | 5 |
| PmKCNQ1 WT–PmKCNE0ΔC90 | *1.9 ± 0.3 | n.d. | n.d. | 5 |
| PmKCNQ1 WT–PmKCNE0ΔC94 | *2.3 ± 0.4 | n.d. | n.d. | 5 |
| PmKCNQ1 WT–PmKCNE0ΔC100 | *1.4 ± 0.2 | n.d. | n.d. | 5 |
| PmKCNQ1 WT–PmKCNE0ΔC105 | 1.9 ± 0.2 | -52.7 ± 0.9 | 2.5 ± 0.1 | 5 |
| PmKCNQ1 WT–PmKCNE0ΔC110 | 1.8 ± 0.1 | -47.5 ± 0.8 | 2.6 ± 0.0 | 5 |
| **Figure 4** | | | | |
| HsKCNQ1 WT | 2.4 ± 0.4 | -26.5 ± 0.8 | 2.5 ± 0.2 | 5 |
| HsKCNQ1 WT–PmKCNE0 WT | 1.6 ± 0.1 | 14.6 ± 1.5 | 1.1 ± 0.1 | 5 |
| DrKCNQ1 WT | 2.4 ± 0.5 | -44.2 ± 1.1 | 1.9 ± 0.0 | 5 |
| DrKCNQ1 WT–PmKCNE0 WT | 0.9 ± 0.0 | -40.4 ± 1.3 | 1.6 ± 0.0 | 5 |
| CiKCNQ1 WT | 1.8 ± 0.3 | -7.2 ± 1.7 | 2.9 ± 0.0 | 5 |
| CiKCNQ1 WT–PmKCNE0 WT | 2.9 ± 0.2 | -10.3 ± 1.1 | 2.6 ± 0.1 | 5 |
| PmKCNQ1 WT–HsKCNE1 WT | 2.7 ± 0.2 | -19.3 ± 1.5 | 1.3 ± 0.0 | 5 |
| PmKCNQ1 WT–HsKCNE3 WT | 2.8 ± 0.2 | -41.3 ± 1.2 | 0.7 ± 0.0 | 5 |
| **Figure 5** | | | | |
| PmKCNQ1 WT–PmKCNE0 H75L | *0.4 ± 0.0 | n.d. | n.d. | 5 |
| PmKCNQ1 WT–PmKCNE0 P73L/F74L/H75L | 0.4 ± 0.0 | -41.3 ± 0.5 | 1.5 ± 0.1 | 5 |
| **Supplementary Figure 5** | | | | |
| PmKCNQ1 WT–PmKCNE0 L54T | *1.8 ± 0.2 | n.d. | n.d. | 5 |

* = normalized to the maximum current amplitude within the recorded region.

| Conditions (VCF) | $I_{max}$ (µA) | $V_{1/2}$ (mV) | *z* | $V_{1/2(F1)}$ (mV) | $V_{1/2(F2)}$ (mV) | *n* |
|---|---|---|---|---|---|---|
| **Figure 2** | | | | | | |
| PmKCNQ1 C205A/G210C | 2.8 ± 0.5 | -87.2 ± 0.8 | 2.0 ± 0.1 | -71.6 ± 1.8 | >80 | 5 |
| PmKCNQ1 C205A/G210C–PmKCNE0 WT | *3.1 ± 0.2 | n.d. | n.d. | -11.8 ± 2.5 | n.d. | 5 |

| Figure | Statistical test | Parameter | Comparisons | P value |
|---|---|---|---|---|
| **Figure 2G** | Tukey–Kramer after one-way ANOVA | G–V $V_{1/2}$ | PmKCNQ1 WT vs LrKCNQ1 WT | 0.0368 |
| | | | PmKCNQ1 WT vs LcKCNQ1 WT | 0.0081 |
| | | | LrKCNQ1 WT vs LcKCNQ1 WT | 0.6849 |
| **Figure 3F** | Tukey–Kramer after one-way ANOVA | G–V $V_{1/2}$ | PmKCNQ1 WT vs PmKCNQ1 WT–PmKCNE0ΔN6 | 0.4405 |
| | | | PmKCNQ1 WT vs PmKCNQ1 WT–PmKCNE0ΔN11 | 0.4082 |
| | | | PmKCNQ1 WT vs PmKCNQ1 WT–PmKCNE0ΔN22 | 0.0415 |
| | | | PmKCNQ1 WT vs PmKCNQ1 WT–PmKCNE0ΔN30 | 0.6950 |
| **Figure 3N** | Tukey–Kramer after one-way ANOVA | G–V $V_{1/2}$ | PmKCNQ1 WT vs PmKCNQ1 WT–PmKCNE0ΔC105 | < 0.001 |
| | | | PmKCNQ1 WT vs PmKCNQ1 WT–PmKCNE0ΔC110 | < 0.001 |
| **Figure 4C** | Unpaired two-tailed Welch's t-test | G–V $V_{1/2}$ | HsKCNQ1 WT vs HsKCNQ1 WT–PmKCNE0 WT | < 0.001 |
| **Figure 4F** | Unpaired two-tailed Welch's t-test | G–V $V_{1/2}$ | DrKCNQ1 WT vs DrKCNQ1 WT–PmKCNE0 WT | 0.0549 |
| **Figure 4I** | Unpaired two-tailed Welch's t-test | G–V $V_{1/2}$ | CiKCNQ1 WT vs CiKCNQ1 WT–PmKCNE0 WT | 0.1607 |
| **Figure 4L** | Unpaired two-tailed Welch's t-test | G–V $V_{1/2}$ | PmKCNQ1 WT vs PmKCNQ1 WT–HsKCNE1 WT | < 0.001 |
| | | | PmKCNQ1 WT vs PmKCNQ1 WT–HsKCNE3 WT | 0.0080 |
