## Supplemental Table 2 for "Characterization of an early-diverging KCNE potassium-channel auxiliary subunit in the jawless vertebrate lamprey"

| Run | Tissue | Total reads before trim (Million reads) | Total reads after trim (Million reads) | % retained after trim (%) | Mean length of R1 before trim (bp) | Mean length of R2 before trim (bp) | Mean length of R1 after trim (bp) | Mean length of R2 after trim (bp) | Q30 bases before trim (%) | Q30 bases after trim (%) |
|---|---|---|---|---|---|---|---|---|---|---|
| SRR9964076 | heart | 266.882 | 254.482 | 95.354 | 150 | 150 | 143 | 143 | 97.8 | 100.0 |
| SRR9964077 | gill | 67.385 | 64.478 | 95.686 | 150 | 150 | 143 | 143 | 98.0 | 100.0 |
| SRR9964078 | testis | 51.819 | 51.030 | 98.478 | 150 | 150 | 148 | 148 | 99.3 | 100.0 |
| SRR9964079 | brain | 286.217 | 273.212 | 95.456 | 150 | 150 | 144 | 143 | 97.9 | 100.0 |
| SRR9964080 | liver | 292.266 | 278.154 | 95.172 | 150 | 150 | 143 | 143 | 97.8 | 100.0 |
| SRR9964081 | oral_gland | 304.972 | 289.959 | 95.077 | 150 | 150 | 144 | 143 | 97.7 | 100.0 |
| SRR9964082 | kidney | 307.687 | 294.386 | 95.677 | 150 | 150 | 144 | 143 | 98.1 | 100.0 |
| SRR9964083 | intestine | 307.983 | 292.761 | 95.057 | 150 | 150 | 144 | 143 | 97.7 | 100.0 |
| SRR9964084 | supraneural_body | 282.527 | 270.428 | 95.717 | 150 | 150 | 144 | 143 | 98.1 | 100.0 |
| SRR9964085 | muscle | 296.558 | 282.051 | 95.108 | 150 | 150 | 144 | 143 | 97.7 | 100.0 |
