## Supplemental Table 3 for "Characterization of an early-diverging KCNE potassium-channel auxiliary subunit in the jawless vertebrate lamprey"

| Run | Tissue | Gene | Number of detected fragments | Total processed fragments | CPM (counts per million of the library) |
|---|---|---|---|---|---|
| SRR9964076 | heart | KCNQ1 | 188 | 127241118 | 1.4775 |
| | | KCNE0 | 190 | 127241118 | 1.4932 |
| SRR9964077 | gill | KCNQ1 | 49 | 32238865 | 1.5199 |
| | | KCNE0 | 50 | 32238865 | 1.5509 |
| SRR9964078 | testis | KCNQ1 | 12 | 25515196 | 0.4703 |
| | | KCNE0 | 0 | 25515196 | 0.0000 |
| SRR9964079 | brain | KCNQ1 | 223 | 136606130 | 1.6324 |
| | | KCNE0 | 183 | 136606130 | 1.3396 |
| SRR9964080 | liver | KCNQ1 | 246 | 139077013 | 1.7688 |
| | | KCNE0 | 224 | 139077013 | 1.6106 |
| SRR9964081 | oral_gland | KCNQ1 | 250 | 144979356 | 1.7244 |
| | | KCNE0 | 245 | 144979356 | 1.6899 |
| SRR9964082 | kidney | KCNQ1 | 224 | 147193056 | 1.5218 |
| | | KCNE0 | 191 | 147193056 | 1.2976 |
| SRR9964083 | intestine | KCNQ1 | 221 | 146380466 | 1.5098 |
| | | KCNE0 | 188 | 146380466 | 1.2843 |
| SRR9964084 | supraneural_body | KCNQ1 | 218 | 135213860 | 1.6123 |
| | | KCNE0 | 180 | 135213860 | 1.3312 |
| SRR9964085 | muscle | KCNQ1 | 224 | 141025603 | 1.5884 |
| | | KCNE0 | 201 | 141025603 | 1.4253 |
